# Phenotypic stasis with genetic divergence

**DOI:** 10.1101/2022.05.28.493856

**Authors:** François Mallard, Luke Noble, Thiago Guzella, Bruno Afonso, Charles F. Baer, Henrique Teotónio

## Abstract

Whether or not genetic divergence on the short-term of tens to hundreds of generations is compatible with phenotypic stasis remains a relatively unexplored problem. We evolved predominantly outcrossing, genetically diverse populations of the nematode *Caenorhabditis elegans* under a constant and homogeneous environment for 240 generations, and followed individual locomotion behavior. Although founders of lab populations show highly diverse locomotion behavior, during lab evolution the component traits of locomotion behavior – defined as the transition rates in activity and direction – did not show divergence from the ancestral population. In contrast, transition rates’ genetic (co)variance structure showed a marked divergence from the ancestral state and differentiation among replicate populations during the final 100 generations and after most adaptation had been achieved. We observe that genetic differentiation is a transient pattern during the loss of genetic variance along phenotypic dimensions under drift during the last 100 generations of lab evolution. These results suggest that short-term stasis of locomotion behavior is maintained because of stabilizing selection, while the genetic structuring of component traits is contingent upon drift history.

## 2 Introduction

Stasis, the lack of directional change in the average values of a trait over time, is the most common phenotypic pattern observed over timespans reaching one million years (Arnold, 2014; Gingerich, 2019; Uyeda et al., 2011). Theory predicts phenotypic stasis when stabilizing selection, or when directional and other forms of selection cancel out over the period examined, acts upon standing genetic variation reflecting the phenotypic effects of mutational input (Charlesworth et al., 1982; Estes and Arnold, 2007; Hansen and Martins, 1996; Lande, 1986; Morrissey and Hadfield, 2012; Stroud et al., 2023). When considering mutation-selection balance on the long-term (as scaled by the effective population sizes), theory has been successfully applied to explain, for example, fly wing evolution over a period of 40 million years (Houle et al., 2017), or nematode embryogenesis over 100 million years (Farhadifar et al., 2015). On the short-term of a few tens to hundreds of generations, however, many natural populations depend on standing genetic variation for adaptation or rescue from extinction, when mutation should be of little influence and founder effects, demographic stochasticity and genetic drift are important (Chelo et al., 2013; Hill, 1982; Mallard et al., 2023a; Matuszewski et al., 2015).

In the short-term, before mutation-selection balance is reached, phenotypic stasis in natural populations is also commonly observed, often despite significant trait heritability and selection (Merilä et al., 2001; Pujol et al., 2018). Explanations for short-term phenotypic stasis have relied on showing that in many cases there were no changes in the breeding traits’ values, that is, no genetic divergence, either because of selection on unmeasured traits that are genetically correlated with observed ones or because of correlated selection due to unknown environmental covariation between observed and unobserved traits with fitness e.g., (Czorlich et al., 2022; Kruuk et al., 2002), both instances of “indirect” selection. Short-term phenotypic stasis without genetic divergence has also been explained by phenotypic plasticity allowing the tracking of environmental fluctuations, e.g., (Biquet et al., 2022; de Villemereuil et al., 2020). Pujol et al. (2018) reviews other processes responsible for phenotypic stasis in the short term. These studies indicate that phenotypic evolution cannot be understood when considering each trait independently of others and that a multivariate description of selection and standing genetic variation is needed. Selection on multiple traits should be seen as a surface with potentially several orthogonal dimensions (Phillips and Arnold, 1989), each with particular gradients depicting selection strength and direction on each trait and between traits (Arnold et al., 2001; Lande and Arnold, 1983). Responses to selection in turn will depend on the size and shape of the **G**-matrix, the additive genetic variance-covariance matrix of multiple traits (Lande, 1979). For example, phenotypic dimensions with more genetic variation are expected to facilitate adaptation, as selection will be more efficient (Lande, 1976, 1979; Schluter, 1996), even if indirect selection can confound predictions about phenotypic evolution (Mallard et al., 2023a; Morrissey and Bonnet, 2019; Stinchcombe et al., 2014).

The extent to which phenotypic stasis is compatible with the expected divergence of the **G**-matrix in the short-term remains little explored cf. (Bohren et al., 1966; Gromko, 1995; Simões et al., 2019; Teotónio et al., 2004; Teotónio and Rose, 2000). Studies in natural populations cannot usually control environmental variation, and estimates of **G**-matrix dynamics are nearly impossible to obtain, while experiments employing truncation selection do not easily model the complexity of the selection surface. Under drift, and assuming an infinitesimal model of trait inheritance, the **G**-matrix size (i.e., the total genetic variance) is reduced and diverges from ancestral states by a factor proportional to the effective population size (Lande, 1976; Lynch and Hill, 1986; Phillips et al., 2001). However, theory that includes the effects of finite population sizes, multivariate selection, and the pleiotropic effects of mutation remains out of reach for changes in genetic covariances between traits and thus **G**-matrix shape (Barton and Turelli, 1987; Burger, 2000; Lande, 1980; Lynch and Walsh, 1998; Simons et al., 2018). We do expect, however, that once most adaptation has occurred, the divergence of the **G**-matrix shape is caused by drift, and also know that different forms of selection might lead to further genetic divergence in the relatively local phenotypic space occupied after adaptation (Doroszuk et al., 2008; Haller and Hendry, 2014). Whether or not genetic divergence will also lead to phenotypic divergence should then depend on the distribution of pleiotropic effects of quantitative trait loci (QTL) alleles, and linkage disequilibrium between them, created by past selection and drift, and ultimately on the developmental and physiological mapping of genetic onto phenotypic variation (Chebib and Guillaume, 2017; Hansen and Wagner, 2001; Morrissey, 2015; Riska, 1989).

Here we seek to find if the short-term evolution of the **G**-matrix follows the directions of selection or if there is loss of genetic variance just by drift. We also seek to determine how genetic divergence is compatible with phenotypic stasis once most adaptation has been achieved. We analyze the evolution of locomotion behavior on the hermaphroditic nematode *Caenorhabditis elegans*, spanning 240 generations of lab evolution in a constant and homogeneous environment, thus maximizing the chances of imposing and detecting stabilizing selection. We could obtain an accurate characterization of the fitness effects of component trait variation of locomotion behavior (transition rates between movement states and direction), by measuring essentially all individuals at the time of reproduction. We expect locomotion behavior to evolve because individual nematodes do not need to engage in foraging and dwelling for feeding (Gray et al., 2005). It is further expected that sexual interaction between hermaphrodites and males impacts the evolution of locomotion behavior (Barr et al., 2018). We characterized the evolution of the broad-sense **G**-matrix for hermaphrodite locomotion behavior, obtained by phenotyping inbred lines derived from the domesticated ancestral population at generation 140 and from three replicate populations during further 50 and 100 generations in the same environment. After domestication, selection gradients were estimated by regressing fertility onto transition rates.

## 3 Methods

### 3.1 Archiving

Data, R code scripts, and modeling results (including **G**-matrix estimates) can be found in our github repository.

### 3.2 Laboratory culture

We analyzed the lab evolution of locomotion behavior during 273 generations (Figure 1A), the first 223 of which have been previously detailed (Noble et al., 2017; Teotónio et al., 2012; Theologidis et al., 2014). Briefly, 16 inbred founders were intercrossed in a 33-generation funnel to obtain a single hybrid population (named A0), from which six population replicates (A[1-6]) were domesticated for 140 generations. Based on the evolution of several life-history traits such as hermaprodite self and outcross fertility, male mating ability or viability until reproduction we have previously shown that most adaptation to lab conditions had occurred by generation 100 (Carvalho et al., 2014a,b; Poullet et al., 2016; Teotónio et al., 2012; Theologidis et al., 2014). From population A6 at generation 140 (A6140), we derived six replicate populations and maintained them in the same environment for another 100 generations (CA[1-6]). CA[1-6] were derived from splitting into six a single pool of at least 10^3^ individuals from large (10^4^) thawed samples of the A6140 population (Theologidis et al., 2014). Inbred lines were generated by selfing hermaphrodites from A6140 (for at least 10 generations), and from CA populations 1-3 at generation 50 and 100 (CA[1-3]50 and CA[1-3]100; Noble et al. (2021)). We refer to these last 100 generations as the focal stage. During the domestication and focal stages, populations were cultured at constant census sizes of *N* = 10^4^ and expected effective population sizes of *N*_*e*_ = 10^3^ (Chelo et al., 2013; Chelo and Teotónio, 2013). Non-overlapping 4-day life-cycles were defined by extracting embryos from plates and seeding starvation-synchronized L1 larvae to fresh food (Teotónio et al., 2012). Periodic storage of samples (> 10^3^ individuals) was done by freezing (Stiernagle, 1999). Revival of ancestral and derived population samples allows us to control for transgenerational environmental effects under “common garden” phenotypic assays (Teotónio et al., 2017).

**Figure 1.**
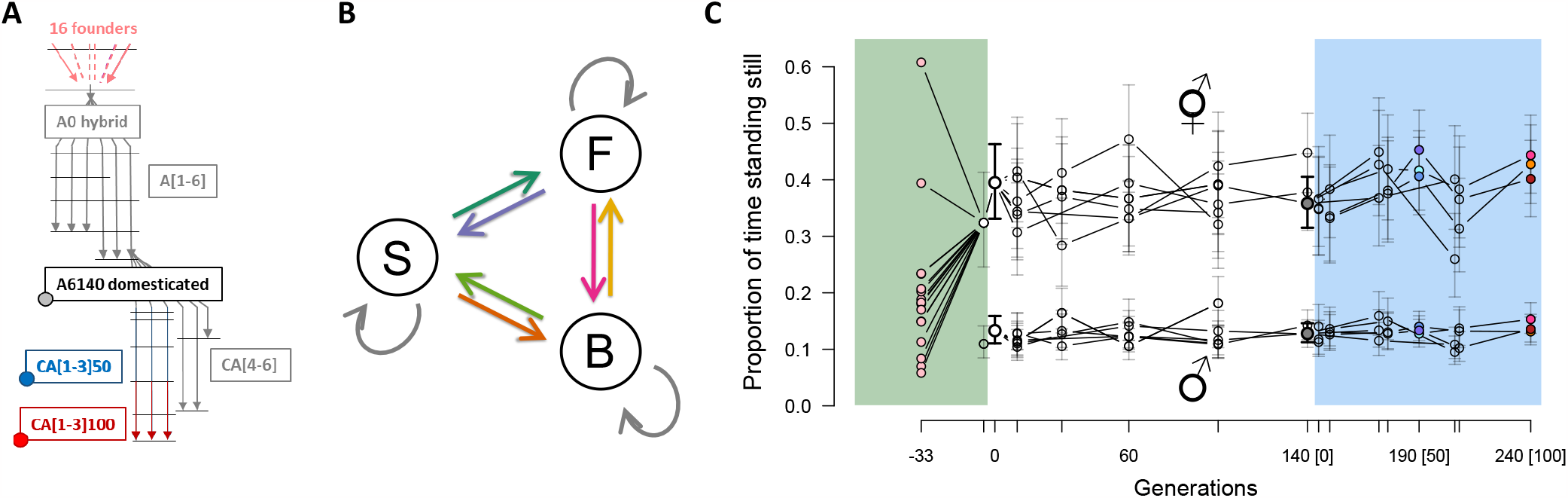
**A**. Experimental design. One hybrid population (A0) was created from the intercross of 16 inbred founders. Six replicate populations were then domesticated to a defined lab environment and after 140 generations one of these (A6140) was the ancestor to six other replicate populations maintained for an extra 100 generations under similar conditions (CA). Inbred lines were derived by selfing hermaphrodites (colored circles) from A6140 and three replicate CA populations at generation 50 and 100 (blue and red). Horizontal lines indicate outbred population samples that were phenotyped. **B**. Modelling locomotion behavior from component traits, defined by the transition rates between moving forward (F), moving backward (B) or being stationary (S). We consider the 6 independent non-self rates, shown in colored arrows. **C**. Evolution of locomotion behavior. Stationary frequency in the founders (pink dots) and outbred populations during lab evolution. Colored overlays indicate three stages of lab evolution: hybridization, domestication and focal. Ticks are sampled time points, while colored points during the focal stage indicate populations from which inbred lines were derived. Point mean estimates are shown for 3-6 replicate populations at other generations, with 95% confidence intervals for each one of them. The evolution of the component traits of locomotion behavior in hermaphrodites and males, the transition rates between movement state and direction, can be found in Figures S4 and S5.

### 3.3 Worm tracking assays

#### 3.3.1 Sampling and design

Population or inbred line samples were thawed from frozen stocks on 9cm Petri dishes and grown until exhaustion of food (*Escherichia coli* HT115). This occurred 2-3 generations after thawing, after which individuals were washed from plates in M9 buffer. Adults were removed by centrifugation, and three plates per line were seeded with 1000 larvae. Samples were maintained for one to two complete generations in the controlled environment of lab evolution. At the assay generation (generation 4-6 post-thaw), adults were phenotyped for locomotion behavior at their usual reproduction time during lab evolution (72h post L1 stage seeding) in single 9 cm plates. At the beginning of each assay we measured ambient temperature and humidity in the imaging room to control for their effects on locomotion. See Table 1 for a detailed description of populations phenotyped with their generations within the different phase of the experiment.

**Table 1:** Identity of the phenotyped populations shown in Figure 1C. The first column presents the population names, composed of a prefix followed by a replicate number. The prefixes “A” and “CA” stand for domestication and focal stages respectively. “CA” populations were all started from the A6 population at generation 140 (A6140) and the generation number restarted at 0. OF5 populations was frozen 5 generations before the A0 during hybridization.

| Population name | Generations | Stage |
| --- | --- | --- |
| OF5, A0 | -5,0 | Ancestral population |
| A1,A2,A3 | 10;30;60;100 | Domestication |
| A4 | 10;30;40;60;100 |  |
| A5,A6 | 10;40;60;100 |  |
| CA1, CA2, CA3 | 5;10;36;50;68;100 | Focal |
| CA4, CA5 | 32;66 |  |
| CA6 | 32 |  |

Inbred lines from the experimental populations were phenotyped over three main common garden experiments in two different lab locations (Lisbon and Paris) by three experimenters. The first common garden included only A6140 lineages, the second CA[1-3]50 lineages and the last one all CA[1-3]100 lineages and A6140 lineages. A6140 G-matrix was initially estimated only from the first common garden (see details below). There were 197 independent thaws, each defining a statistical block containing 2-22 samples. 188 inbred lines from the A6140 population were phenotyped, with 52 CA150, 52 CA250, 51 CA350, 51 CA1100, 53 CA2100 and 68 from CA3100 (not including the A6140 lineages from the third common garden). Each line was phenotyped in at least two blocks (technical replicates). CA[1-3]50 and CA[1-3]100 lines were phenotyped within a year. A6140 lines were phenotyped over two consecutive years. A set of 63 A6140 lineages that were phenotyped together with the CA[1-3]100 populations in the third common garden were used to compute a second A6140 G-matrix. We further phenotyped the outbred populations and the 16 founders in a single common garden. For these, there were 9 independent thaws, of which 5 also contained founders. All founders and populations were phenotyped twice except for A6140, which was included in six blocks.

To improve the estimation of the selection surface in our lab evolution environment (see below), we also assayed locomotion bias in 56 inbred lines derived from populations evolved in a high-salt environment (GA[1,2,4]50) for which fertility data was available (Noble et al., 2017). These lines were phenotyped in the same blocks as the A6140 lines included in the gamma matrix analysis (first common garden, single experimenter). Removing these lines from the analysis did not affect the mode of the posterior distribution estimates of our coefficients. It only led to the loss of statistical power reflected by wider credible intervals (analysis not shown).

#### 3.3.2 Imaging

To measure locomotion behavior we imaged adults 72h post-L1 seeding using the Multi-Worm Tracker [MWT version 1.3.0; Swierczek et al. (2011)]. Movies were obtained with a Dalsa Falcon 4M30 CCD camera and National Instruments PCIe-1427 CameraLink card, imaging through a 0.13-0.16 mm cover glass placed in the plate lid, illuminated by a Schott A08926 backlight. Plates were imaged for approximately 20-25 minutes with default MWT acquisition parameters. Choreography was used to filter and extract the number and persistence of tracked objects and assign movement states across consecutive frames as forward, still or backwards, assuming that the dominant direction of movement in each track is forward (Swierczek et al., 2011).

MWT detects and loses objects over time as individual worms enter and leave the field of view or collide with each other. Each track is a period of continuous observation for a single object (the mapping between individual worms and tracks is not 1:1). We ignored the first 5 minutes of recording, as worms are perturbed by plate handling. Each movie contains around 1000 tracks with a mean duration of about 1 minute. The MWT directly exports measurements at a frequency that can vary over time (depending on tracked object density and computer resource availability), so data were standardized by subsampling to a common frame rate of 4 Hz. Worm density, taken as the mean number of tracks recorded at each time point averaged over the total movie duration, was used as a covariate in the estimation of genetic variance-covariances below.

### 3.3.3 Differentiating males from hermaphrodites

A6140 and all CA populations are androdioecious, with hermaphrodites and males segregating at intermediate frequencies (Teotónio et al., 2012; Theologidis et al., 2014). We were able to reliably (97% accuracy) differentiate between the sexes based on behavioral and morphological traits extracted from MWT data.

We first evaluated a set of simple descriptions of individual size, shape, and movement to find a subset of metrics that maximized the difference in preference for a two-component model between negative and positive controls: respectively, inbred founders and two monoecious (M) populations which contained no, or very few, males; and three dioecious (D) populations with approximately 50% males [M and D populations were derived from A6140, see Theologidis et al. (2014) and Guzella et al. (2018)]. Starting with worm area, length, width, curvature, velocity, acceleration, and movement run length as parent traits from the Choreography output, derived descendant traits were defined by first splitting parents by individual movement state (forward, backward, still) and calculating the median and variance of the distribution for each track. Traits with more than 1% missing data were excluded, and values were log-transformed where strongly non-normal (a difference in Shapiro-Wilk −*log*_10_(*p*) > 10). Fixed block and log plate density effects were removed by linear regression before fitting the residuals to two-component Gaussian mixture models. These two-component Gaussian models were fit to tracks for each line/population [R package *mclust* Scrucca et al. (2016), *VII* spherical model with varying volume], orienting labels by area (assuming males are smaller than hermaphrodites). We sampled over sets of three traits, requiring three different parent trait classes, at least one related to size. We took the set maximizing the difference in median Integrated Complete-data Likelihood (ICL) between control groups (log area, log width, and velocity, all in the forward state). By this ranking, the 16 inbred founders and two monoecious populations fell within the lower 19 samples (of 77), while the three dioecious populations fell within the top 15 samples.

To build a more sensitive classifier robust to male variation beyond the range seen in control data, we then trained an extreme gradient boosting model using the full set of 30 derived traits on the top/bottom 20 samples ranked by ICL in the three-trait mixture model [R package *xgboost*, Chen and Guestrin (2016)]. Negative control samples were assumed to be 100% hermaphrodite, while tracks in positive controls were assigned based on *mclust* model prediction, excluding those with classification uncertainty in the top decile. Tracks were classified by logistic regression, weighting samples inversely by size, with the best cross-validated model achieving an area under the precision-recall curve of 99.75% and a test classification error of 3.1% (*max depth* = 4, *eta* = 0.3, *subsample* = 0.8, *eval metric* = “*error*”). Prediction probabilities were discretized at 0.5.

Males tend to move much faster than hermaphrodites (Lipton et al., 2004), and because individual collision leads to loss of tracking, sex is strongly confounded with track length and number. To estimate male frequencies at the sample level, tracks were sampled at 1s slices every 30s over each movie in the interval 400-1200 seconds, and line/population estimates were obtained from a binomial generalized linear model (Venables and Ripley, 2002). Estimates appear to saturate at around 45%, presumably due to density-dependent aggregation of multiple males attempting to copulate.

### 3.4 Locomotion behavior

#### 3.4.1 Definition of transition rates

In a one-dimensional space, individual locomotion behavior can be described by the transition rates of activity and direction. We modeled the expected sex-specific transition rates between forward, still and backward movement states with a continuous time Markov process. We consider a system having *d* = 3 states with *P*(*t*_1_, *t*_2_) ∈ ℜ^*d,d*^, *t*_2_ > *t*_1_, denoting the transition probability matrix (Jackson, 2011; Kalbfleisch and Lawless, 1985):

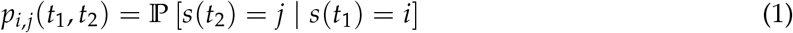

where *s*(*t*) ∈ *t*, with *t* = {*still, <u>f</u>orward, <u>b</u>ackward*} being the movement state occupied in instant *t*. We consider a time-homogeneous process described by the transition rate matrix:

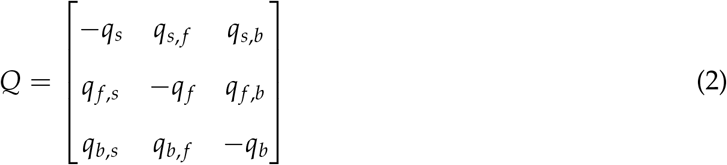

where *q*_*i,j*_ ≥ 0 ∀*i, j*, subject to the constraint:

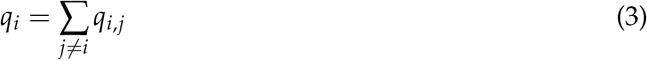

Hence, six of the nine possible transitions are independent. Let *θ* denote the parameters to be estimated, containing the off-diagonal elements from equation 2:

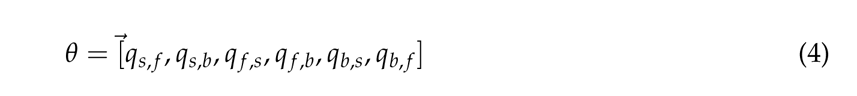

In this model, an object’s time remains in a given state is on average 1/*q*_*i*_. Since the process is stationary, the probability of transition is a function of the time difference Δ*t* = *t*_2_ − *t*_1_, such that *P*(*t*_1_, *t*_2_) = *P*(Δ*t*), and the elements of the *P*(Δ*t*) matrix:

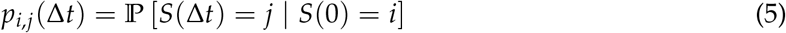

It then follows that:

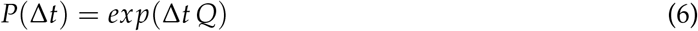

where exp(·) denotes the matrix exponential. The constraint in equation 3 ensures that:

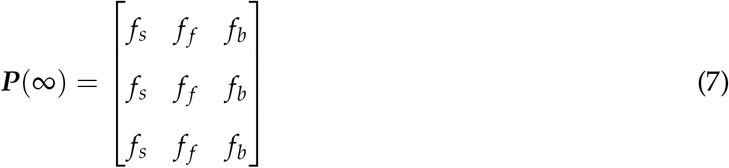

where *f*_*i*_ is the relative frequency of state *i* that no longer depends on the previous state (all three rows of the ***P***(∞) matrix converge). We find that the state frequencies from ***P*(∞)** are a monotonic and mostly linear function of the observed frequencies of movement states (Figure S3), showing that violations of the Markov assumption of the model do not induce a large bias in the long-term predictions of our model.

#### 3.4.2 Estimation of transition rates

To estimate transition rates, we have *N* objects (individual tracks) from each technical replicate (Petri plate), with the data on the *k*-th object denoted as:

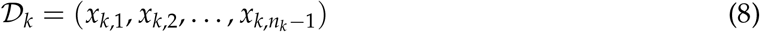

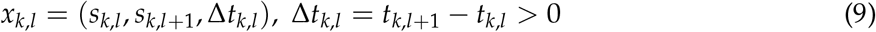

where *s*_*k,l*_ is the state of the *k*-th object in the *l*-th time-point in which it was observed, and *t*_*k,l*_ is the instant of time in which this observation was made. Then, given data *D* = {*D*_1_, *D*_2_, …, *D*_*N*_}, the log-likelihood for the model for analysis is (Bladt and Sorensen, 2005; Kalbfleisch and Lawless, 1985):

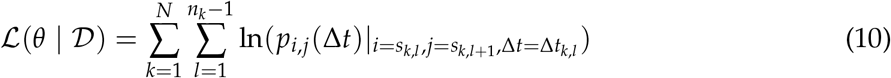

where *p*_*i,j*_ (Δ*t*) was defined in equation 5, and is calculated as a function of the parameters *θ* via equation 4. Therefore, the data on the *N* objects can be represented as the number of observations of *x* = (*i, j*, Δ*t*), which we denote as *ñ*_*i,j*,Δ*t*_ :

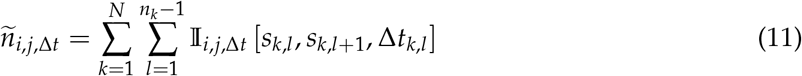

and where I_*i,j*,Δ*t*_ [·] is the indicator function:

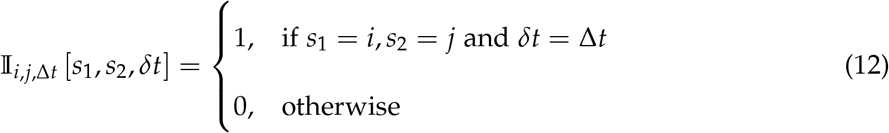

The input data can then be compressed by considering only the data:

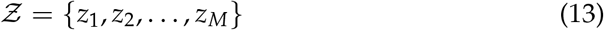

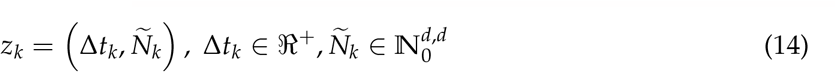

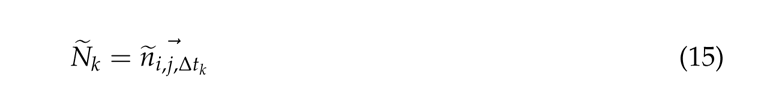

The log-likelihood to estimate transition rates can be finally rewritten as:

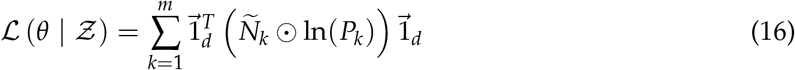

where 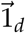 is a *d*-dimensional vector of 1s, ⊙ denotes the Hadamard product, and ln *P*_*k*_ is the matrix obtained by taking the logarithm of each value in matrix *P*_*k*_.

These models were specified using RStan (Stan Development Team (2018), R version 3.3.2, RStan version 2.15.1), which performs Bayesian inference using a Hamiltonian Monte Carlo sampling to calculate the posterior probability of the parameters given the observed data. We used multi-log normal prior distributions with mean transition rate and a coefficient of variation: ln(*q*_*i,j*_) ∼ *N*(ln(2), 0.6).

Throughout, we denote non-self transition rates *q*_*k*_ the six off-diagonal elements of the **Q** matrix estimated by the above model.

#### 3.4.3 Male and inbreeding effects

Using the transition rates measured in populations and inbred lines, we fit a series of linear mixed-effects models to test for phenotypic evolution in the outbred populations, for effects of male frequency on hermaphrodite transition rates in the outbred populations, and for inbreeding effects in the inbred lines. Given sparse temporal sampling, we make the conservative assumption of independence of observations within domestication and focal stages. For transition rate *q*_*k*_:

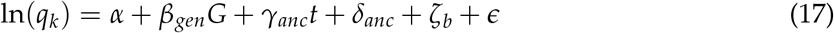

with *α* the trait mean, *β*_*gen*_ a fixed effect of generation number *t, γ*_*anc*_ and *δ*_*anc*_ random effects accounting for intercept and slope differences between the domestication and focal periods of lab evolution (both∼ *N*(0, *σ*^2^)), *ζ* ∼ *N*(0, *σ*^2^) a random effect of block *b* and *ϵ* ∼ *N*(0, *σ*^2^) the residual error.

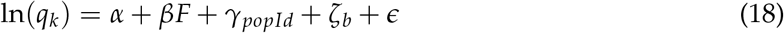

with *β* a fixed effect of male frequency F, *γ*_*popId*_ ∼ *N*(0, *σ*^2^) a random effect accounting for differences between populations, *ζ* ∼ *N*(0, *σ*^2^) a random effect of block *b*, and *ϵ* ∼ *N*(0, *σ*^2^) the residual error.

As we estimate the **G**-matrix from the line differences (see next section), it is likely that it does not reflect the true additive genetic (co)variance matrix (**G**-matrix) unless the mean trait values among lines are similar to the mean trait values of the outbred population from which the lines were derived (Lynch and Walsh, 1998). Only with directional, genome-wide, dominance or epistasis would the “broad-sense” **G**-matrix not be a good surrogate for the “narrow-sense” additive **G**-matrix. See Chapter 3 of Kearsey and Pooni (1996) for the different ways dominance and epistasis can change segregation variance in F2 crossing designs. Because the lines and the populations were phenotyped at different times, we included environmental covariates:

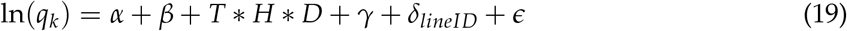

where environmental covariates: temperature (**T**), relative humidity (**H**) and density (**D**) are fitted as fixed effects. *β* is a two-level categorical fixed effect (inbred lines or population). *γ* is a two-level categorical fixed effect accounting for differences between the years of phenotyping measurements of the A6140 lineages. *δ* ∼ *N*(0, *σ*^2^) a random effect accounting for line identity within populations and *ϵ* ∼ *N*(0, *σ*^2^) the residual error.

Both male and inbreeding models were fit using the *lmer* function in R package *lme*4, and non-zero values of fixed effects were tested against null models without fixed effects with likelihood ratio tests. Marginal *r*^2^ for the male frequencies were computed using the *r*.*squaredGLMM* function of the package *MuMIn* (Bartoń, 2020).

### 3.5 Transition rate genetics

#### 3.5.1 G-matrix estimation

Genetic (co)variances of transition rates per population are estimated as half the between inbred line differences for lines separately derived from the evolving outbred populations. In the absence of selection during inbreeding and canceling of directional non-additive gene action, this broad-sense **G**-matrix obtained from inbred lines is an adequate surrogate for the additive **G**-matrix of outbreeding populations (Kearsey and Pooni, 1996; Lynch and Walsh, 1998). We test these assumptions (see below).

**G**-matrices for the six non-self transition rates *q*_*k*_ were estimated from trait values for the inbred lines derived from focal populations. We estimated **G**-matrices separately for each of the seven populations (A6140, CA[1-3]50, CA[1-3]100). The 6 transition rates *q*_*k*_ were fitted as a multivariate response variable *y* in the model:

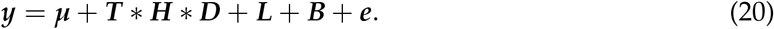

where the intercept (***μ***) and the environmental covariates: temperature (**T**), relative humidity (**H**) and density (**D**) were fitted as fixed effects. Environmental covariates were fitted individually and with all possible interactions. Each covariate was standardized to a mean of 0 and standard deviation of 1. Block effects (**B**) and line identities (**L**) were modeled as random effects and **e** was the residual variance. We then estimated a matrix of genetic (co)variance as half the line covariance matrix (**L**). An additional two-level categorical effect was included when estimating the A6140 matrix that accounts for differences between the 2012 and 2013 phenotyping blocks in the first common garden. As mentioned above, a second A6140 matrix was computed from the data collected in the third common garden using the same model.

For modeling we use the R package *MCMCglmm* (Hadfield, 2010). We constructed priors as the matrix of phenotypic variances for each trait. Model convergence was verified by visual inspection of the posterior distributions and by ensuring that the autocorrelation remained below 0.05. We used 100000 burn-in iterations, a thinning interval of 2000 and a total of 2100000 MCMC iterations.

#### 3.5.2 G-matrices under random sampling

For each of our seven populations (A6140, CA[1-3]50, CA[1-3]100), we constructed 1000 randomised **G**-matrices to generate a null distribution against which to compare the observed estimates. We randomly shuffled both the inbred line and block identities and fit equation 20. We then computed the posterior means of our 1000 models to construct a null distribution.

We additionally generated 1000 matrices for the A6140 population using the same procedure on random subsets of 60 (of 188 total) inbred lines to determine the effects of sampling the same number of lines as those for CA[1-3]50 and CA[1-3]100 populations.

#### 3.5.3 G-matrix divergence and differentiation

We define genetic divergence whenever comparing derived from ancestral **G**-matrices, and similarly, genetic differentiation when comparing the **G**-matrices of replicate populations at generation 50 or generation 100.

To compare the overall size of the **G**-matrices during experimental evolution, we first computed the trace of the matrices and then performed spectral analyses of the posterior ancestral **G**-matrix. The decomposition of the posterior ancestral **G**-matrix allows one to describe the overall **G**-matrix shape, with the relative genetic variance between the six eigenvalues of each eigenvector, indicating whether the matrix is elliptical (a few large eigenvalues) or round (homogeneous eigenvalues). The first eigenvector (defined as *g*_*max*_) is the linear combination of traits where the genetic variance is maximized.

To compare changes in **G**-matrix shape, we computed the angles Θ between the eigenvectors of the **G**-matrices as in Noble et al. (2019). For *g*_*i*_ and *g*_*j*_ two eigenvectors:

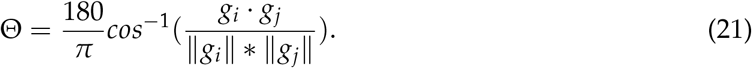

As both *g*_*i*_ (*g*_*j*_) and −*g*_*i*_ (-*g*_*j*_) are eigenvectors of the same *G*-matrix, Θ values between 90° and 180° were transformed so that Θ always remains between 0° and 90°. We sampled the posterior distribution of the *G*-matrix of interest for each angle to create a credible interval. The null expectation for Θ is calculated as the angle between 1,000 pairs of random vectors sampled from a uniform distribution *U*^6^(−1, 1) as in Mallard et al. (2023a).

In our final analysis of genetic divergence and differentiation, we used eigentensor methods to explore differences between the **G**-matrices, following (Aguirre et al., 2014; Hine et al., 2009).

Genetic (co)variance tensors (**Σ**) are fourth-order objects describing how phenotypic dimensions between transition rates maximize differences between all the **G**-matrices. The genetic variation among multiple **G**-matrices can be described by **Σ** decomposition into orthogonal eigentensors (*E*_*i*_, with *i* being the orthogonal dimensions), each associated with an eigenvalue quantifying its contribution to variation in **Σ** (***α***_***i***_). In turn, eigentensors can be decomposed into eigenvectors (*e*_*ii*_), each with associated eigenvalues (*λ*_*i*_). Aguirre et al. (2014) implemented this approach in a Bayesian framework using *MCMCglmm*, and Morrissey and Bonnet (2019) made an important modification to account for sampling where the amount of variance in ***α***_***i***_ is compared to an expected distribution by sampling a finite number of lines.

### 3.6 Selection on transition rates

#### 3.6.1 Selection surface

The log-transformed, covariate-adjusted fertility values (best linear unbiased estimates) for each inbred line were downloaded from Noble et al. (2017), exponentiated, and divided by the mean to obtain a relative fitness measure (*w*_*l*_).

Since we did not observe any directional change in locomotion behavior or component transition rates during lab evolution, and because the inbred lines were derived after domestication, most of adaptation to the lab environments has occurred, and we do not expect linear (directional) selection to be significant (but see below). We estimated quadratic selection gradients using partial regression, following (Lande and Arnold, 1983), with the *MCMCglmm* R package:

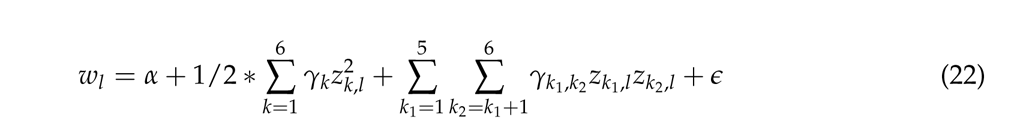

with *α* being the mean relative fitness among all lines and *γ* the partial coefficients estimating quadratic selection on each transition rate *k*, or between pairs of transition rates *k*_1_ and *k*_2_. Environmental covariates (temperature, humidity, density) were defined and normalized as for the **G**-matrices estimation described above. Model residuals were normal and homocedastic (not shown).

We compared the results of this model (equation 22) with those of linear mixed effect models including as a random effect the additive genetic similarity matrix **A** between inbred lines, as defined in Noble et al. (2017) and Noble et al. (2021). We have also compared results from equation 22 to models including coefficients for linear selection on each transition rate. Under both circumstances parameter estimates are similar to those presented, albeit with changing credible intervals (not shown). Including other measured traits by the worm tracker, such as body size [a trait related to developmental time that is known to affect fertility in our populations (Theologidis et al., 2014)] similarly does not affect the qualitative conclusions we reach.

#### 3.6.2 G-matrix alignment with the selection surface

We used canonical analysis (Phillips and Arnold, 1989) to visualize the selection surface as:

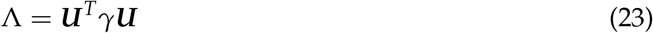

with **U** being the matrix of eigenvectors of ***γ***, and Λ the diagonal matrix of eigenvalues (denoted *λ*_[1−6]_). **G**-matrices were rotated to visualize them as:

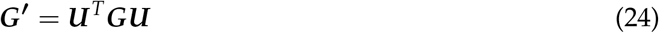

To sample a null distribution of the *γ* eigenvalues along the rotated dimensions, we fit the same model after permuting the relative fitness values of the lines. We then extracted the diagonal elements of these permuted *γ* after rotation using the estimated **U**.

To see the evolution of the **G**-matrix in the selection surface, we calculated the Pearson product moment correlations between the eigentensor vectors explaining most of the genetic differences between the 7 matrices (*e*_11_, *e*_12_) with the canonical selection dimensions (*y*_1_-*y*_6_). We estimated uncertainty in these values by sampling from the posterior distribution of *γ* 1000 times.

### 3.7 Inference of effects

Most of our analysis relies on Bayesian inference of genetic or phenotypic effects. As discussed in Walter et al. (2018), the “significance” of effects can be inferred when there is no overlap between the posterior null sampling distributions with the posterior empirical estimate of the expected values. Thus, we compare expected value estimates such as a mean or mode with the 95% credible intervals under random sampling of the expected value. The “significance” of the posterior mode estimates is based on their overlap with the posterior null distribution of the posterior modes (Walter et al., 2018). For all comparisons of posterior distributions significance can be inferred when their 83% credible intervals do not overlap (Austin and Hux, 2002), assuming homoscedasticity.

## 4 Results

### 4.1 Laboratory culture

Our lab evolution system is based on a hybrid population derived from 16 founder strains (Figure 1A). Replicate samples from the hybrid population were domesticated for 140 non-overlapping generations at census size *N*=10^4^ to an environment in part characterized by constant density, temperature and relative humidity, and by little spatial structure during the life-cycle (see Methods). The dynamics of several life-history traits during domestication indicate that most adaptation to lab conditions occurred by generation 100 (Carvalho et al., 2014a,b; Poullet et al., 2016; Teotónio et al., 2012; Theologidis et al., 2014). From a single domesticated population we derived replicate populations and evolved them for another 100 generations in the same environmental conditions. Although we measured locomotion behavior throughout of lab evolution, we only follow the **G**-matrix of its component traits during the last 100 generations, after adaptation, a stage that we call here the focal stage of lab evolution (Figure 1A).

*C. elegans* reproduces mostly by selfing in nature though there is considerable variance in male mating performance among the founders (Teotónio et al., 2006). By training a model on a suite of size- and locomotion-related metrics, we found that hermaphrodites could be clearly differentiated from males (see Methods), and estimated males frequencies were high during the entire experiment (Figure S1). Because *C. elegans* are androdioecious, and hermaphrodites cannot mate with each other, average expected selfing rates at a generation are 1 minus twice the male frequency at the previous generation (Teotónio et al., 2012), and we can conclude that outcrossing was the predominant reproduction mode during lab evolution. Previously, we showed that effective population sizes during domestication were of about *N*_*e*_=10^3^ (Chelo and Teotónio, 2013).

### 4.2 Evolution of locomotion behavior

We measured locomotion behavior at the time of reproduction for each outbred population and the inbred founders using worm video tracking (Swierczek et al., 2011). The output, after quality control and initial analysis, are individual worm tracks categorized at a given point in time by activity (moving, or not) and direction (forwards or backwards). We model a three-state memoryless (Markov) process with homogeneous spatial and temporal dynamics (see Methods, Figure S2). We view this as an obviously false but useful approximation of worm locomotion behavior under our conditions, which is only partially violated (worms tend to resume forward movement more often than expected; Figure S3). Component traits of locomotion behavior are the (sex-specific) six non-self transition rates between forward movement, backward movement, and immobility.

We find that while the founders of lab evolution show great diversity in locomotion behavior under lab conditions, evolved populations rapidly attained, and maintained, a stable level after hybridization for 240 generations. For example, considering the proportion of time individual worms are stationary (Figure 1B), we observe values of around 40% for hermaphrodites - much higher than most founders - while males are much more vagile (stationary around 10%). Neither hermaphrodite nor male transition rates showed a directional change from the hybrid ancestral state over the full 240-generation period (Table 2, Supplementary Figures S4 and S5). Differences between replicate populations can be explained by sampling error.

**Table 2:** Phenotypic stasis: Results of anova LRT 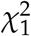 tests for directional changes in mean transition rates in hermaphrodites (H) and males (M), during the 240 generations of lab evolution. Corrected P values for multiple comparisons were obtained with the Benjamini-Hochberg method. Transition rates notation XY stands for transition from trait X to Y, S: Still, F: Forward and B: Backward.

| Transition rate | Sex | Chisq | P value | P adjusted |
| --- | --- | --- | --- | --- |
| SF | H | 0.542 | 0.462 | 0.799 |
| SB | H | 0.715 | 0.398 | 0.799 |
| FS | H | 0.389 | 0.533 | 0.799 |
| FB | H | 3.077 | 0.079 | 0.799 |
| BS | H | 0.037 | 0.847 | 0.924 |
| BF | H | 1.810 | 0.179 | 0.799 |
| SF | M | 0.414 | 0.520 | 0.799 |
| SB | M | 0.647 | 0.421 | 0.799 |
| FS | M | 0.542 | 0.462 | 0.799 |
| FB | M | 0.053 | 0.817 | 0.924 |
| BS | M | 0.001 | 0.980 | 0.980 |
| BF | M | 0.082 | 0.774 | 0.924 |

### 4.3 Broad-sense **G**-matrix

To estimate **G**-matrices, we used approximately 200 lines from the generation 140 domesticated population (A6140), and approximately 50 lines from each of three replicate populations derived from A6140 and sampled at generations 50 (CA[1-3]50) and 100 (CA[1-3]100) of the focal lab evolution. We use these broad-sense **G**-matrices as a surrogate for the narrow-sense (additive) **G**-matrices of the outbred populations (see Methods). These two kinds of matrices might not be identical because of selection during inbreeding or because of differential expression of non-additive genetic effects in inbred and outbred individuals. Such differences, if present, manifest as differences in the mean values of inbred and outbred samples as directional effects will statistically average out for polygenic traits (Kearsey and Pooni, 1996; Lynch and Walsh, 1998). We used the inbred lines and the focal A6140 ancestor to compare means for all transitions and we did not find any evidence of directional non-additive genetic effects (Table 3).

**Table 3:** Inbreeding effects: Results of anova LTR 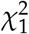 testing for mean phenotypic differences between the mean of the inbred lines and the mean of the A6140 population from which they were derived. Corrected P values for multiple comparisons were obtained with the Benjamini-Hochberg method. Transition rates notation XY stands for transition from trait X to Y, S: Still, F: Forward and B: Backward.

| Transition rates | Chisq | P values | P adjusted |
| --- | --- | --- | --- |
| SF | 5.445 | 0.020 | 0.118 |
| SB | 1.319 | 0.251 | 0.419 |
| FS | 0.107 | 0.743 | 0.860 |
| FB | 1.443 | 0.230 | 0.419 |
| BS | 1.170 | 0.279 | 0.419 |
| BF | 0.031 | 0.860 | 0.860 |

Our **G**-matrices could also differ from the **G**-matrices of outbred populations due to the absence of males in the inbred lines; which were abundant in the outbred populations. This is because males are known to disturb hermaphrodite locomotion behavior (Lipton et al., 2004). We tested for effects of male frequency on transition rates in outbred populations with univariate linear models and found that they were weak at best (Figure S6).

### 4.4 Genetic divergence and differentiation

For the domesticated 140 population (A6140), ancestral to all CA populations during further 100 generations in the same environment after adaptation, there is significant genetic variance in all hermaphrodite transition rates, relative to a null distribution from permutations of line and technical replicate identity (Figure 2A). Likewise, the posterior distributions of most (12 of 15) covariance estimates between transition rates do not overlap 0, and differ from the null distribution of posterior means. The A6140 **G**-matrix is structured in two main behavioral modules, with the transitions from still to forward or backward (i.e. leaving the still state) showing positive covariances with each other and negative covariances with other transition rates.

**Figure 2.**
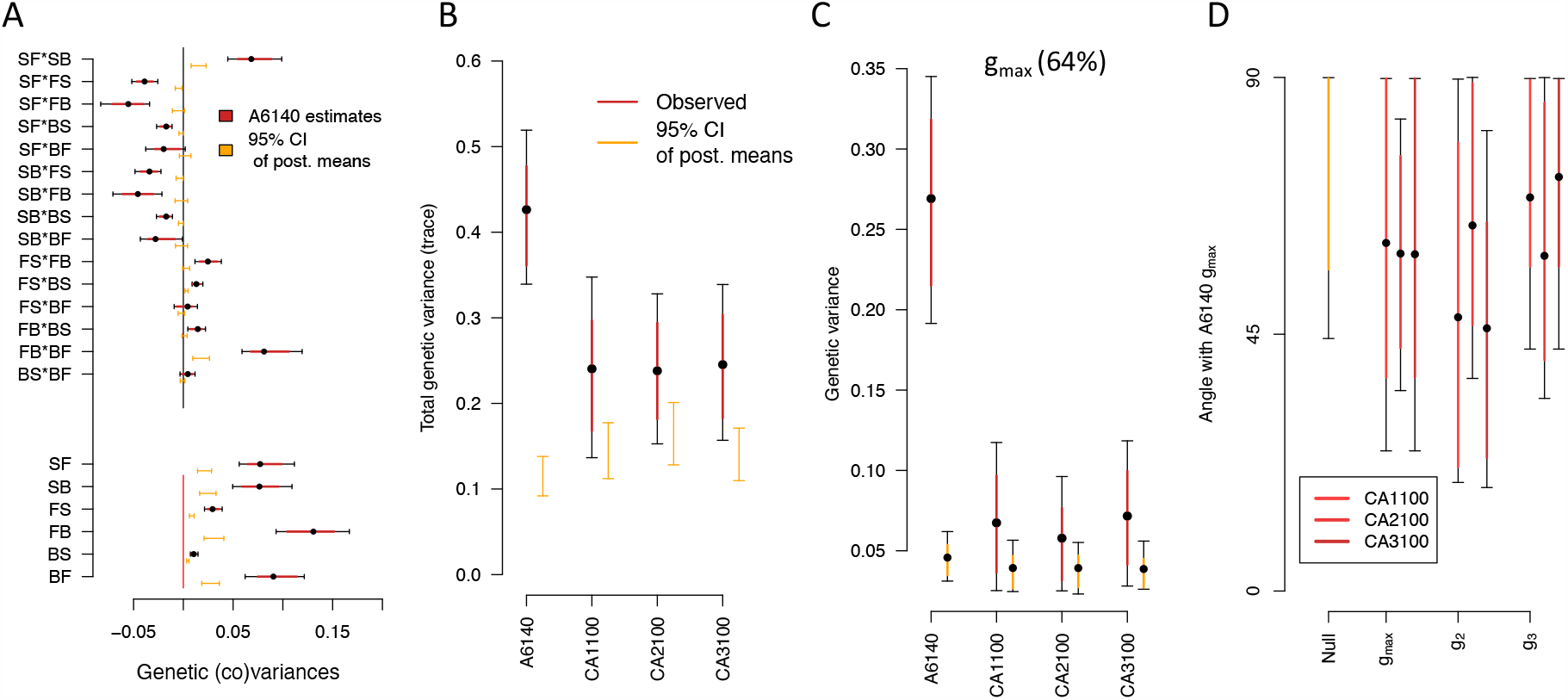
**G**-matrix evolution and divergence during the focal stage. **A**. A6140 **G**-matrix. Shown are the 15 genetic covariances between transitions rates (top) and six the genetic variances of transition rates (bottom), as bars and dots the 95% and 83% credible intervals (black and red) and mean of the posterior distribution, respectively. “S”, “F”, “B” stand for still, forward and backward movement states, with letter ordering indicating the direction of movement. **G**-matrices of the CA populations can be found in Supplementary Figure S7. **B**. Total amount of genetic variance computed as the sum of the **G** diagonal elements (trace). All observed posterior means differ from the null 95% of posterior means (orange). **C**. Genetic variance along the phenotypic dimension encompassing most genetic variation in the ancestral A6140 population (*g*_*max*_, red mean, 83% and 95% CI). A6140 *g*_*max*_ explains 64% of the total genetic variance Table 4). D. The angle (Θ, see Methods) between the A6140 *g*_*max*_ and the first three eigenvectors of the evolved **G**-matrices (*g*_*max*_, *g*_2_ and *g*_3_ of the CA[1-3]100 populations). There is no alignment between the evolved populations’ first three eigenvectors with the ancestral *g*_*max*_. Dots show the mean estimate with bars the 83% and 95% credible interval of the posterior **G**-matrix distribution. The null expectation was obtained by computing the angle between pairs of random vectors sampled from a uniform distribution.

Inbred lines from the ancestral and evolved populations at generations 50 and 100 were phenotyped in separate common garden assays. CA[1-3]50 inbred lines show a clear difference relative to other generations in all transition rate variances and mean body area or velocity (not shown). We thus phenotyped a subset of A6140 lines together with all CA[1-3]100 lines and ensured that these measurements were comparable (see Figure S9). As a consequence, we only compare **G**-matrix between A6140 and CA[1-3]100 populations, though we discuss the divergence among the three CA[1-3]50 **G**-matrices as they were phenotyped in a single common garden assay.

When looking at the evolved CA populations, we see that their **G**-matrices are reduced after 100 generations of evolution (Figure S7). Reduced genetic (co)variance in generation 100 is particularly obvious when calculating the trace of the **G**-matrices, although all populations contain more genetic variance than expected by chance (Figure 2B). The loss of genetic (co)variances during focal evolution could be due to differences in statistical power or the result of continued lab evolution. Sub-sampling A6140 to the sampling sizes of CA[1-3]100 populations, while increasing the credible intervals did not affect the estimated modes, with many of them remaining different from the null (Figure S8). This difference is robust to common garden assay variation (see Figure S9A).

Eigendecomposition of the A6140 **G**-matrix further shows that, for the phenotypic dimension encompassing 64% of genetic variation in this population (*g*_*max*_), the projected variance of CA[1-3]100 populations in this dimension is much reduced (Figure 2C). In this *g*_*max*_ dimension of maximal ancestral variation, leaving the still movement states (still-to-forward, and still-to-backwards, transition rates) are positively associated with each other while being negatively associated with all other transition rates (Table 4). Therefore, the size of the **G**-matrix has evolved over 100 generations.

**Table 4:** Eigendecomposition of the A6140 G-matrix. Transition rates notation XY stands for transition from trait X to Y, S: Still, F: Forward and B: Backward.

|  | **A6140 population** |  |  |  |  |  |
| --- | --- | --- | --- | --- | --- | --- |
| | $g_{\max}$ | $g_2$ | $g_3$ | $g_4$ | $g_5$ | $g_6$ |
| Eigenvalues | 0.263 | 0.105 | 0.022 | 0.01 | 0.005 | 0.003 |
| HPD lower | 0.196 | 0.074 | 0.016 | 0.007 | 0.003 | 0.002 |
| HPD upper | 0.348 | 0.156 | 0.03 | 0.014 | 0.006 | 0.003 |
| Proportion | 0.645 | 0.257 | 0.054 | 0.025 | 0.012 | 0.007 |
| *Trait loadings:* |  |  |  |  |  |  |
| SF | -0.438 | 0.474 | 0.086 | 0.286 | -0.539 | -0.451 |
| SB | -0.423 | 0.444 | -0.488 | 0.198 | 0.501 | 0.31 |
| FS | 0.214 | -0.315 | -0.176 | 0.717 | 0.297 | -0.472 |
| FB | 0.629 | 0.383 | -0.596 | -0.162 | -0.236 | -0.143 |
| BS | 0.112 | -0.127 | -0.074 | 0.533 | -0.488 | 0.666 |
| BF | 0.419 | 0.563 | 0.602 | 0.234 | 0.276 | 0.119 |

Next, we performed eigendecomposition of the CA[1-3]100 and compared if the shape of the evolved **G**-matrices aligned with the *g*_*max*_ of the ancestral population. We find that the angle (Θ, see Methods) between the A6140 *g*_*max*_ and the first three eigenvectors of the evolved **G**-matrices (*g*_*max*_, *g*_2_ and *g*_3_) are much higher than zero, and indeed are not different from random expectations (Figure 2D). At generation 50, however, only one replicate population diverged in **G**-matrix shape because the angle between A6140 *g*_*max*_ and the CA[1,3]50 *g*_*max*_ is smaller than expected by chance (Figure S11) – note that although the matrices were estimated in different common garden assays, the direction of the canonical dimensions can be compared –. Overall, the size and the shape of the **G**-matrix evolved over 100 generations.

We more formally tested for divergence of the **G**-matrices from the ancestral state during 100 generations and for differentiation between derived replicate populations at generation 50 using eigentensor analysis (see Methods). This analysis identifies the phenotypic dimensions along which there are most differences between the several matrices being compared.

When looking for divergence between A6140 and CA[1-3]100, the first eigentensor, ***E***_**1**_, explains more variation than the null expectation (Figure 3A, 73%). **G**-matrix coordinates in the space of ***E***_**1**_ (Figure 3B), show that the A6140 population drives most significant differences between all matrices, and thus encompasses most of the genetic divergence. Along the first eigenvector of ***E***_**1**_ (called *e*_11_;Figure 3C), divergence is mostly due to the loss of genetic variance in the CA populations (Figure 2C). Confirming this interpretation, we further find that *e*_11_ is highly collinear with the *g*_*max*_ of the A6140 population, the phenotypic dimension encompassing most ancestral genetic variation (not shown). Similar conclusions can be reached when comparing the A6140 results from the third common garden with the CA[1-3]100 populations while ensuring that the assay period does not affect the variance estimates (see Figure S9). The first three eigenvectors of CA[1-3]100 populations did not align with the A6140 *g*_*max*_ (Figure 2D), though by generation 50 CA[1,3]50 replicates were aligned (Figure S11). Together, these findings indicate that the genetic variance is progressively dispersed over multiple phenotypic dimensions during focal evolution.

**Figure 3.**
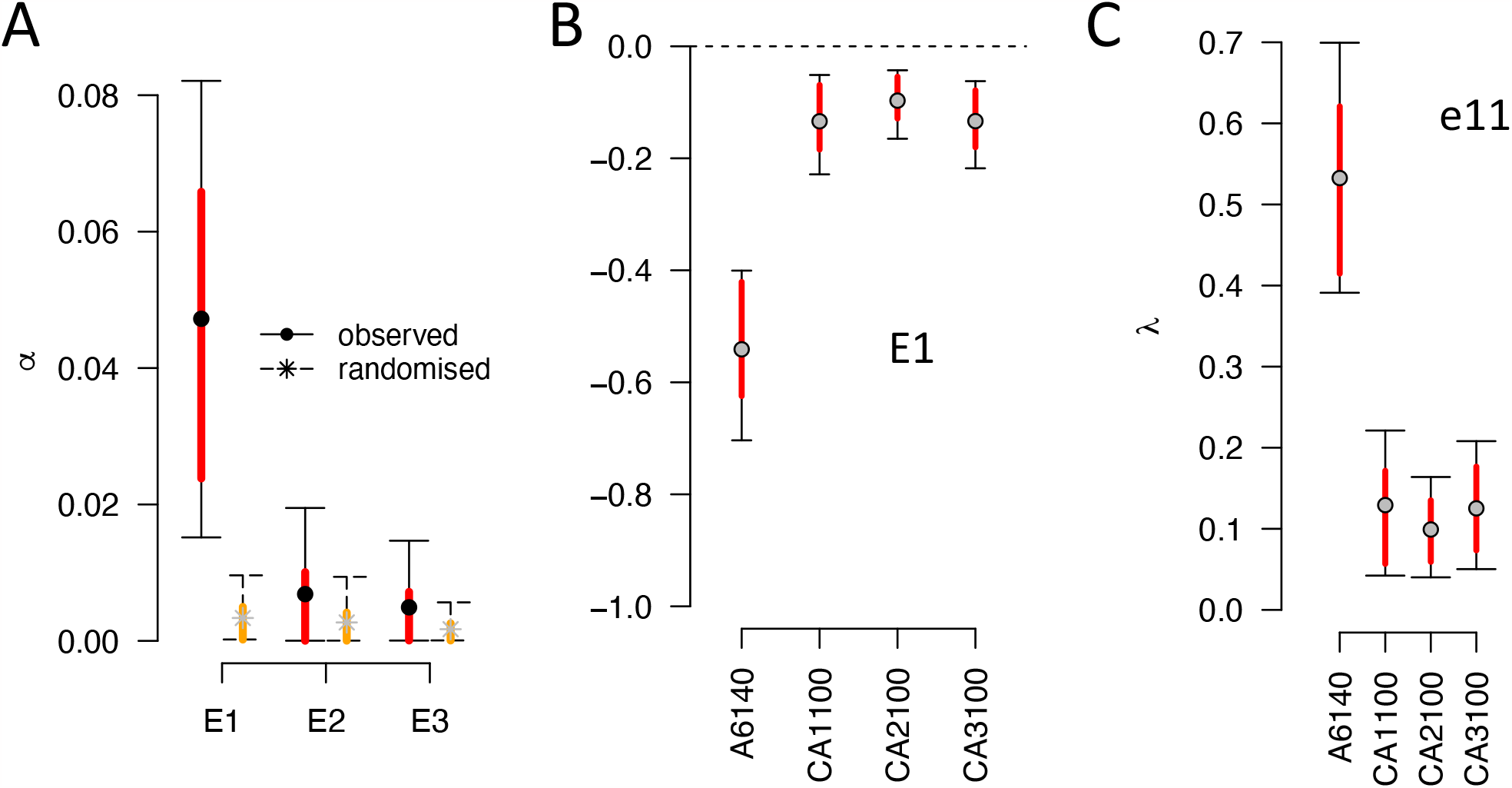
Genetic divergence. **A**. Spectral decomposition of variation among A6140 and CA[1-3]100 **G**-matrices. The variance ***α***_***i***_ associated with the *i*th eigentensor ***E***_***i***_ is compared to a null permutation model where variation among matrices is due to sampling (see Methods). Although several eigentensors are different from zero (black bars, 95% credible interval) only the first one, ***E***_**1**_, do not overlap the null (red and orange bars, 83% credible intervals). **B**. The coordinates of the **G**-matrices in the space of the first eigentensor ***E***_**1**_ for each population tested. Absolute values of the coordinates in the first eigentensor represents its contribution to the difference between matrices. **C**. Contribution of specific transition rate combinations to coordinated changes among **G**-matrices. The amount of genetic variance in the direction of the greatest variation among all **G**s for the first eigenvector of ***E***_**1**_ (*e*_11_), for each population. Eigentensor decomposition of the CA[1-3]50 **G**-matrices, testing for differentiation at generation 50, can be found in Figure S9.

We tested for differentiation between replicate populations during focal evolution by restricting the spectral analysis to only the three CA[1-3] **G**-matrices, separately at generation 50 and generation 100. For the CA[1-3]50 populations, we observe that a single eigentensor was different from the null expectations, explaining 53% of the differences between the three **G**-matrices (Figure S10). The coordinates of these matrices in the space of the eigentensor indicate that CA150 and the remaining two populations contributed in opposite directions to the difference observed. Most of this difference is expressed along the first two eigenvectors (50% and 37%): CA[2-3]50 lost variance along the first eigenvector and CA150 along the second one. A similar analysis at generation 100 did not show differentiation between the three CA[1-3]100 **G**-matrices (not shown).

### 4.5 Selection on locomotion behavior

In Noble et al. (2017) we reported the fertility of many inbred lines used to estimate the **G**-matrices. This data encompasses hermaphrodite self-fecundity and progeny viability until early larvae, measured in an environment that closely mimicked that of lab evolution. With this data at hand we can estimate the selection surface of locomotion in our lab environment by applying equation 22, with relative fertility being partially regressed onto the transition rates (see Methods).

We find that the 95% credible intervals for several coefficients for correlated selection between pairs of transition rates do not overlap zero: negative between still-forward (SF) and forward-still (FS) and positive between SB and FS, and FS and BS (Figure S12). To visualize the selection surface, we rotated the *γ*-matrix with canonical analysis (see Methods). The resulting selection surface suggests a saddle with three unstable equilibria in three canonical dimensions *y*_1_-*y*_3_, indicating disruptive selection, and three stable equilibria in three dimensions (*y*_4_-*y*_6_), indicating stabilizing selection (Figure 4, Table 5). We only find, however, evidence of weak and strong stabilizing selection at *y*_5_ and *y*_6_, respectively, because only these empirical estimates are unlikely under the null distribution.

**Table 5:**
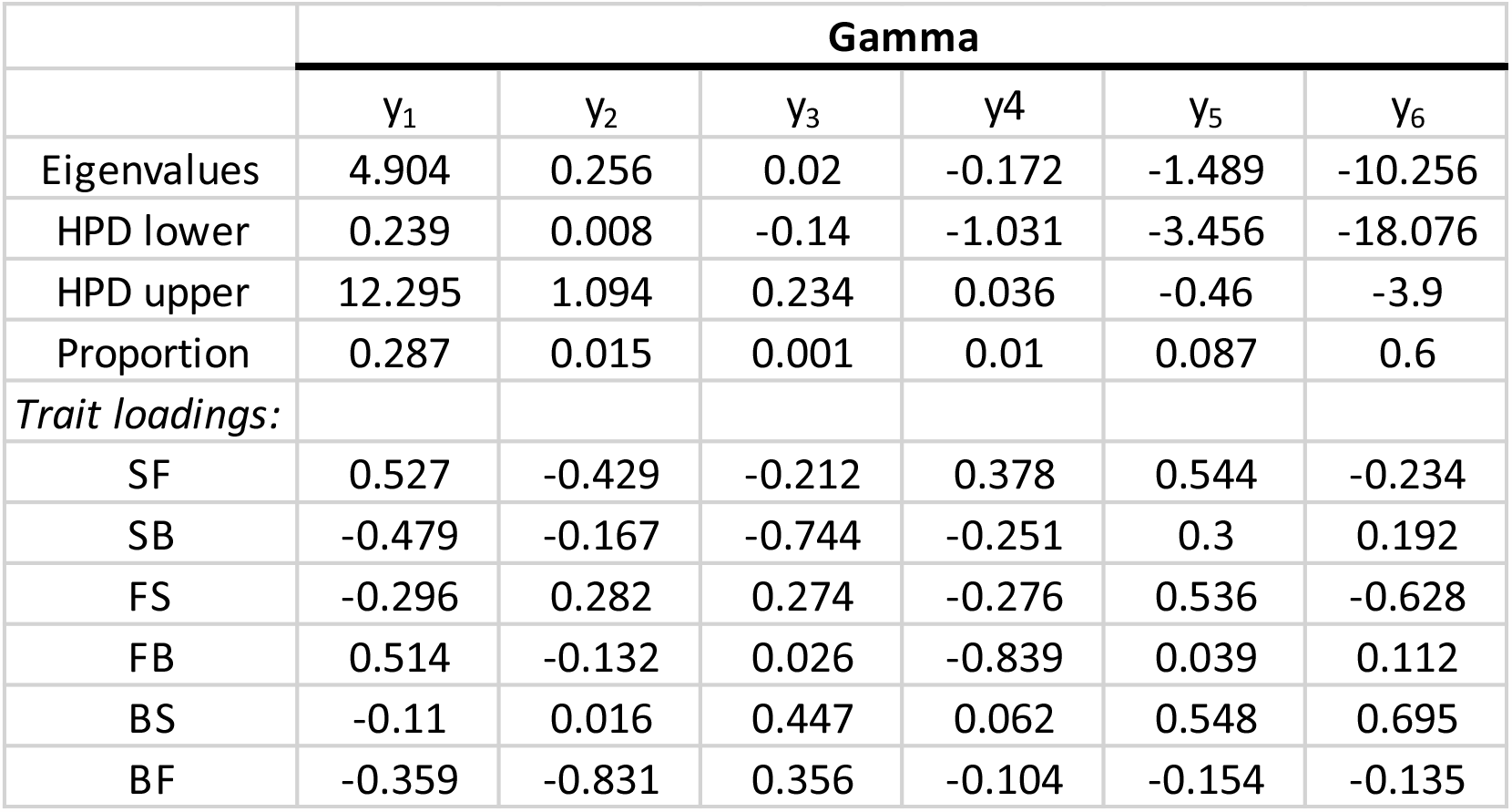
Eigendecomposition of the *γ* G-matrix. Transition rates notation XY stands for transition from trait X to Y, S: Still, F: Forward and B: Backward.

|  | **Gamma** |  |  |  |  |  |
| --- | --- | --- | --- | --- | --- | --- |
| | $\gamma_1$ | $\gamma_2$ | $\gamma_3$ | $\gamma_4$ | $\gamma_5$ | $\gamma_6$ |
| Eigenvalues | 4.904 | 0.256 | 0.02 | -0.172 | -1.489 | -10.256 |
| HPD lower | 0.239 | 0.008 | -0.14 | -1.031 | -3.456 | -18.076 |
| HPD upper | 12.295 | 1.094 | 0.234 | 0.036 | -0.46 | -3.9 |
| Proportion | 0.287 | 0.015 | 0.001 | 0.01 | 0.087 | 0.6 |
| *Trait loadings:* |  |  |  |  |  |  |
| SF | 0.527 | -0.429 | -0.212 | 0.378 | 0.544 | -0.234 |
| SB | -0.479 | -0.167 | -0.744 | -0.251 | 0.3 | 0.192 |
| FS | -0.296 | 0.282 | 0.274 | -0.276 | 0.536 | -0.628 |
| FB | 0.514 | -0.132 | 0.026 | -0.839 | 0.039 | 0.112 |
| BS | -0.11 | 0.016 | 0.447 | 0.062 | 0.548 | 0.695 |
| BF | -0.359 | -0.831 | 0.356 | -0.104 | -0.154 | -0.135 |

**Figure 4.**
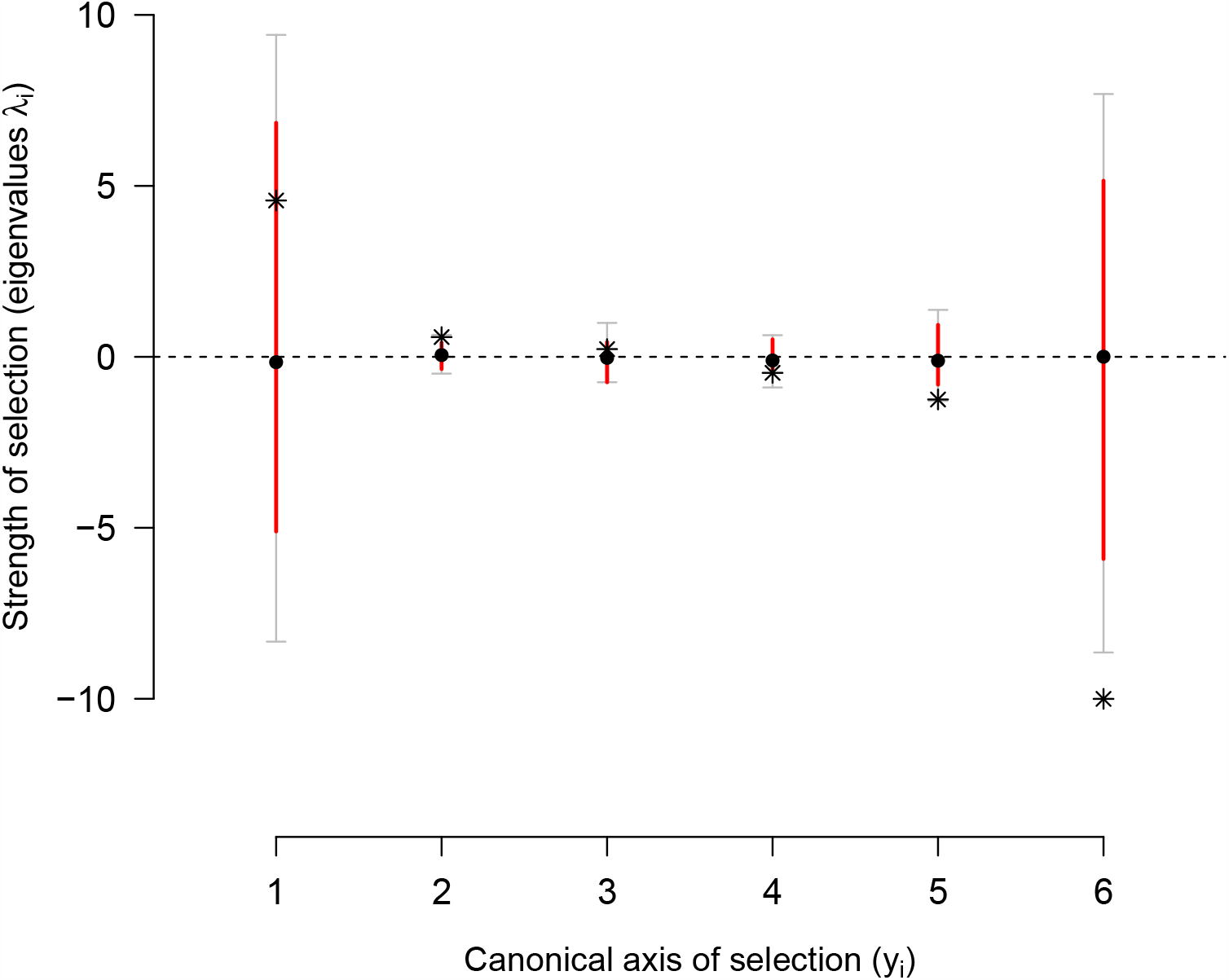
Selection surface of locomotion behavior. Canonical analysis of the *γ*-matrix shows positive phenotypic dimensions (*y*_1_-*y*_3_) of transition rate combinations under disruptive selection (as measured by the eigenvalue *λ*), and negative dimension (*y*_4_-*y*_6_) under stabilizing selection. Stars show the mode of the posterior empirical distribution (see Methods). These estimates are to be compared to the posterior distribution of null modes (dots and colored bars, the mean and 83% and 95% credible intervals). The *γ*-matrix before canonical rotation can be found in Figure S12.

### 4.6 G-matrix evolution in the selection surface

Projection of the **G**-matrices onto the canonical selection dimensions shows that most genetic variance is concentrated in dimensions (*y*_2_-*y*_4_), while the dimensions under stabilizing selection (*y*_5_ and *y*_6_) do not show much genetic variance that can be lost after generation 140 (Figure 5). *y*_1_ does similarly not show much genetic variance. Along all potential selection dimensions, loss of genetic variance is consistent with drift when assuming an infinitesimal model of trait inheritance (Barton et al., 2017) and effective population sizes of *N*_*e*_ = 10^3^ (Chelo and Teotónio, 2013). However, for the *y*_1_ and *y*_6_ dimensions, initial and evolved populations at generation 100 clearly varied less than the founders’ isolates of experimental evolution, as their 83% posterior distributions do not overlap.

**Figure 5.**
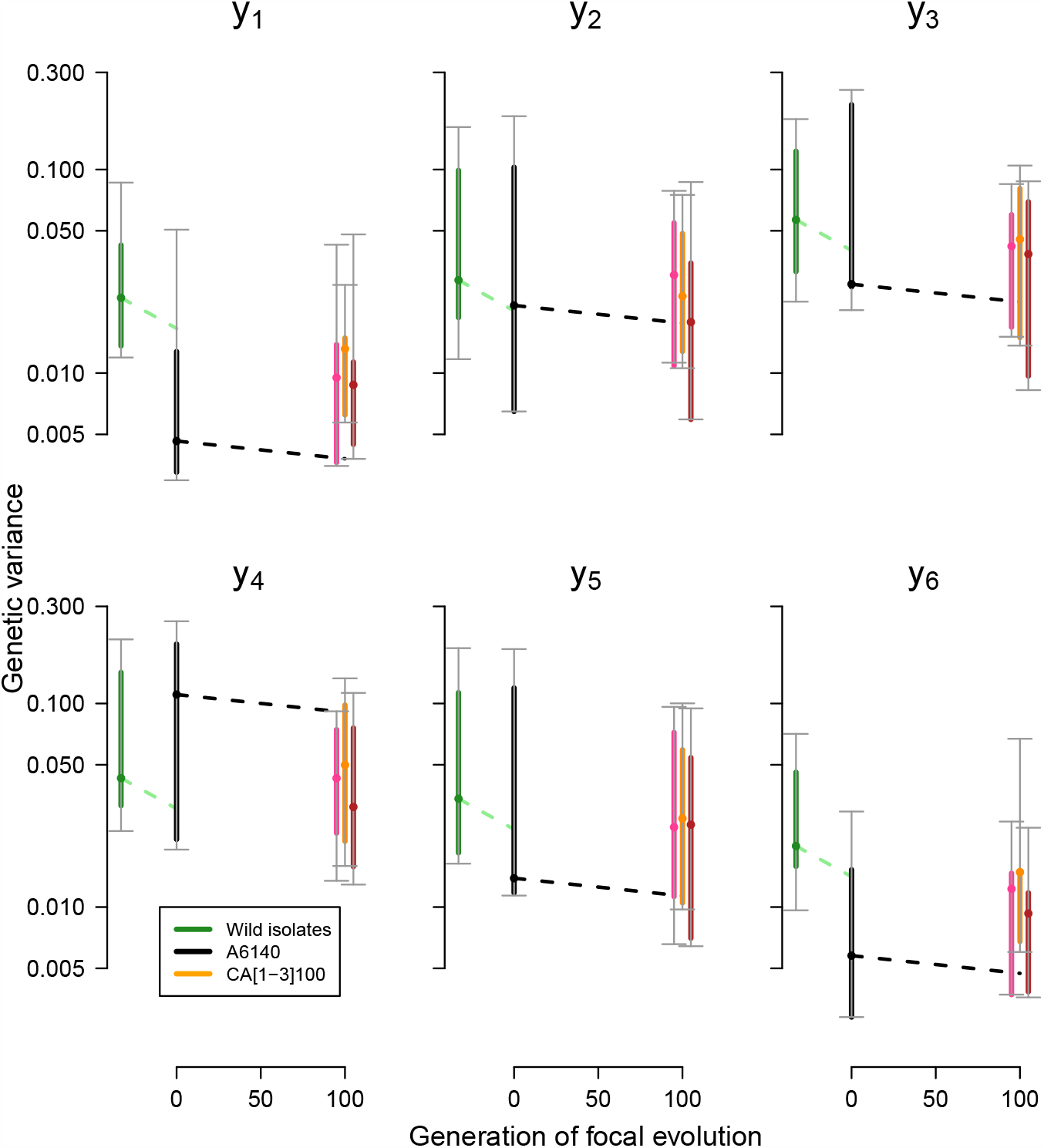
**G**-matrix evolution in the selection surface. Loss of genetic variance along axes *y*_2_-*y*_5_, which contain most of the genetic variance in the evolved populations and are under very weak or no selection, is compatible with expectations from genetic drift under the assumption of infinitesimal trait inheritance (dashed lines, for *N*_*e*_ = 10^3^). We modelled drift as a per generation loss of genetic variance of 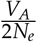. Along *y*_1_ and *y*_6_, genetic variance was much reduced relative to the founders of experimental evolution (green). The genetic variance of each canonical axis *y*_*i*_ was obtained by rotation of the original **G**-matrices, with 95% (grey) and 83% (colored) credible intervals from sampling 400 matrices in the posterior distributions for each **G**-matrix. Dots show the median estimates. See also Figure Figure S13.

To assess if **G**-matrix evolution aligned with the selection surface, we calculated the correlation between the directions of genetic divergence at generation 100 of the focal stage (Figure 3), and differentiation of replicate population at generation 50 (Figure S10), with the canonical selection dimensions (Figure 4). Overall there is a strong alignment of both genetic divergence and genetic differentiation axes obtained from our two tensor analysis with *y*_3_ (Figure S13), meaning that drift fully explains genetic divergence and differentiation.

## 5 Discussion

The evolution of *C. elegans* locomotion behavior during 240 generations in a fairly constant and homogeneous lab environment is characterized by stasis, following a genetically and phenotypically dynamic 33-generation period of hybridizing the founder strains (Figure 1). We observed that most genetic variance along the several phenotypic dimensions was reduced during domestication and before the focal stage (Figure 5), presumably because of stabilizing selection. After generation 140, during the focal stage, genetic variance continued to be lost (Figure 2), though not obviously along the phenotypic dimensions where stabilizing selection was detected (Figure 4, Figure 5). Most, if not all, of the genetic divergence (Figure 2, Figure 3, Figure S11) and of the genetic differentiation among generation 50 populations (Figure S10) is sufficiently explained by genetic drift (Figure 5, Figure S13). Despite divergence and differentiation being sufficiently explained by genetic drift, there was stabilizing selection after domestication because the inbred lines used for selection estimates were from the focal stage (Figure 4, Figure S12). Although we did not formally test for directional selection, it is likely that it only occurred during the hybridization of the founders (Figure 1, Figure 5). We further suspect that the main phenotypic dimensions under stabilizing selection (*y*_5_ and *y*_6_, Figure 4) lost genetic variance during this initial hybridization stage. Stabilizing selection and genetic drift therefore appear to have maintained phenotypic stasis over the available phenotypic variation after founder hybridization. A future area of research would be to ask if specific mechanisms of density and/or frequency-dependent selection – such as sex allocation, sexual selection, viability selection during early larval growth, maternal effects, etc.; as we before described in some of the same populations (Carvalho et al., 2014b; Chelo et al., 2013; Dey et al., 2016; Poullet et al., 2016) –, underlie stabilizing selection on locomotion behavior. It would further be interesting to test if starting lab evolution from founders whose average locomotion behavior is away from the phenotypic space measured in our populations converge into a similar “adaptive zone” cf. (Simpson, 1944; Uyeda et al., 2011), or continue to be somehow constrained by standing genetic variation.

It can be argued that, with only 16 founders, we have little power to reject the hypothesis that there was no loss and that the genetic (co)variances we found after domestication reflect natural standing genetic variation. At mutation-drift balance, the **G**-matrix should reflect the patterns of mutational effects described by the **M** matrix, the equivalent measure of trait mutational variances, and covariances between them due to pleiotropy (Lande, 1979; Lynch and Hill, 1986). Elsewhere, we have estimated the **M** matrix in two of the founders of lab evolution, which show locomotion values divergent from those of lab evolution populations, by phenotyping a set of about 120 lines that accumulated mutations in a nearly neutral fashion for 250 generations (Baer et al., 2005; Yeh et al., 2017). We found that the **M** matrices from these founders have similar sizes and are well aligned with each other, but not with the **G**-matrix of our A6140 domesticated population (Mallard et al., 2023b). Loss of genetic variances from the founders during hybridization was, therefore, partly because of directional selection. Future work should nonetheless try to understand if mutation-selection balance is responsible for the maintenance of genetic variation in locomotion behavior in nature by comparing **G**-matrices from natural populations, as they can be obtained from a large collection of wild isolates now available (Cook et al., 2017; Lee et al., 2021), with **M** matrices (Houle et al., 1996; Johnson and Barton, 2005).

One of the major findings is that of divergence and the transient differentiation of the **G**-matrix during the last 100 generations of lab evolution. The phenotypic dimensions of genetic divergence and differentiation among all populations were not aligned with the phenotypic dimensions under selection, and most, if not all, of the genetic variance lost during this focal 100-generation period was expected with drift. Not unexpectedly, loss of genetic variance mostly occurred along the dimensions with most genetic variance in the ancestral lab-adapted population (*g*_*max*_). This dimension may represent a continuum between activity and direction of movement in foraging and dwelling, expressed by the positive association between transition rates from the still state (Flavell et al., 2020; Gray et al., 2005). Stabilizing selection favors a negative association between transition rates from the still state, which, elsewhere, we have shown is under directional selection in a new stressful environment (Mallard et al., 2023a). As was the case here, however, transition rates from the still state in the new stressful environment did not evolve under directional selection because of a lack of relevant genetic variation in the appropriate direction. Overall, these observations are congruent with experiments in *D. melanogaster* by K. Fowler and colleagues where, after bottlenecking an outbred population, there was a reduction in the size of the **G**-matrix for wing morphology in the derived bottlenecked populations, and size divergence among them, as expected under drift (Fowler and Whitlock, 1999; Phillips et al., 2001). Genetic differentiation also occurred because the shape of the **G**-matrix changed as derived populations showed different genetic covariances between traits. Interestingly, drift history was consequential to the future phenotypic divergence of particular bottlenecked populations in a new environment (Whitlock et al., 2002). We suspect a similar result would have been observed had we performed experimental evolution in a new environment and had the differentiated replicates from generation 50 of the focal stage as ancestral populations.

Most of our analyses and the underlying theoretical predictions are predicated on the assumption that the infinitesimal model of trait inheritance is a good approximation of the truth. However, that assumption may be violated, as the genetic variances and covariances of locomotion behavior will not in the short-term of our hybridization and lab evolution be independent of allele frequency changes and linkage disequilibrium between smaller effect quantitative trait loci (QTL). QTL allele frequency independence is expected only in the long-term when approaching strong recombination and weak selection, mutation and drift, steady-states (Barton, 1990; Barton et al., 2017; Vladar and Barton, 2014). Our findings pose the question of how genetic drift, together with stabilizing selection, generates variable allele frequency changes at QTL so that pleiotropy or linkage disequilibrium between them eventually results in genetic covariances that diverge from the ancestral states and are not common among replicate populations. Even if populations eventually lose most genetic variance, this transient differentiation could be important for future phenotypic evolution if the environment changes. In our case, recombination during the focal stage should have remained much weaker than selection between 0.5-1 cM regions (Chelo and Teotónio, 2013; Noble et al., 2017, 2021); for total a total genome size of 300 cM. If after domestication several QTL alleles within these linked regions segregate at low frequency, it is possible that selection and drift was such that each replicate population during divergence fixed alleles with differently signed phenotypic effects that would not average out when comparing across populations (Bernstein et al., 2019; Cohan, 1984; Gromko, 1995). Inflation of the effects of drift is further expected because there is a correlation across generations between the traits’ breeding values of successful parents and their offspring, resulting in a reduction in effective population sizes (Robertson, 1961; Santiago and Caballero, 1998).

Short-term phenotypic stasis without genetic divergence in natural populations has been explained by indirect selection or phenotypic plasticity, among several other processes cf. (Estes and Arnold, 2007; Pujol et al., 2018), despite heritability and direct selection on the traits that were followed. Our study shows that phenotypic stasis can also occur with simultaneous genetic divergence and transient genetic differentiation. We conclude that the adaptive landscape in our lab environment is best understood as a table-top mountain, where a plateau with potentially very shallow optima are of no consequence to genetic or phenotypic divergence. In the long term, phenotypic stasis is a common pattern observed over up to a million years. For longer periods, rapid divergence in mean trait values is observed from fossil records, or inferred from phylogenetic trees, potentially because new adaptive zones are accessible after extreme ecological changes. Given our results, we speculate that upon such changes, phenotypic divergence and differentiation of populations can be facilitated by cryptic evolution of genetic covariance structure during phenotypic stasis.

## 6 Acknowledgments

We thank A. Crist, J. Garcia, H. Gendrot, C. Goy, V. Pereira, F. Melo, and A. Silva for help with worm handling and data acquisition; R. Costa, R. Kerr, and N. Scwierczek for help with hardware and software implementation; N. Barton, C. Dillmann, A. Futschik, L. Kollar, S. McDaniel, P. Phillips, S. Proulx, A. Le Rouzic, and A. Veber for discussion. We also thank F. Guillaume, B. Pujol and P. Simões for suggestions that improved the presentation of this work.

## 8 Author contributions

Conceptualization FM, LN, CB, HT; hardware and software implementation BA, TG; data acquisition and analysis BA, FM, LN, TG; funding acquisition LN, CB, HT; project administration HT; resources CB, HT; writing, original draft FM, HT; writing, review and editing LN, CB; correspondence FM and HT.

## 10 Supplementary Figures

**Figure S1:**
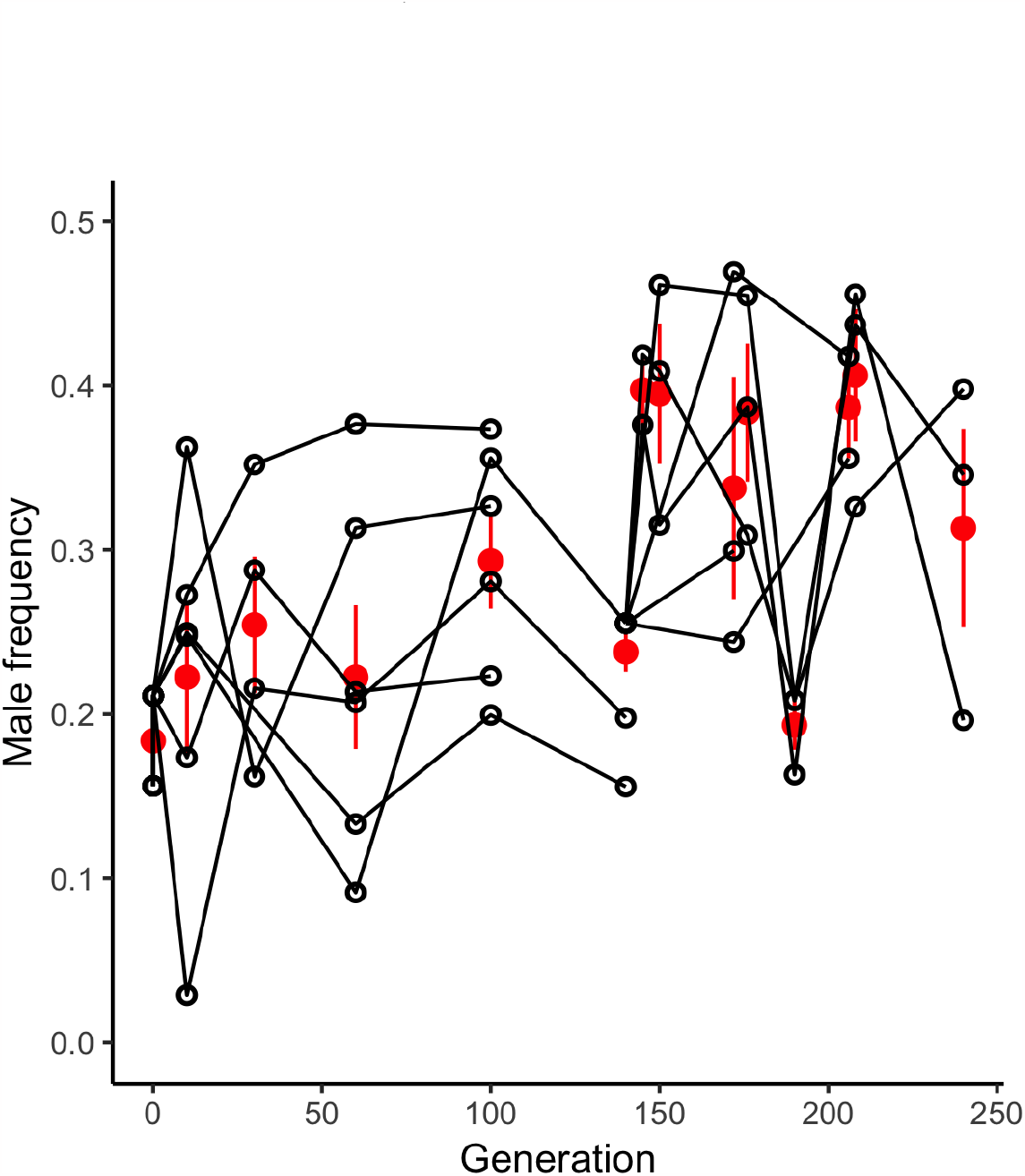
Male frequencies during lab evolution. Males and hermaphrodite tracks were differentiated with a 30-trait classifier based on moments of size, shape and velocity-related traits derived from Multi-Worm Tracker metrics, and frequencies were estimated from 1s slices across movies. Empty circles indicate the estimates for each replicate population (between 1 and 6 at each time point), red circles the mean among replicate populations (± standard error). During the first 100 generations of domestication, the estimates are similar to those obtained by directly counting the number of males (Teotónio et al., 2012).

**Figure S2:**
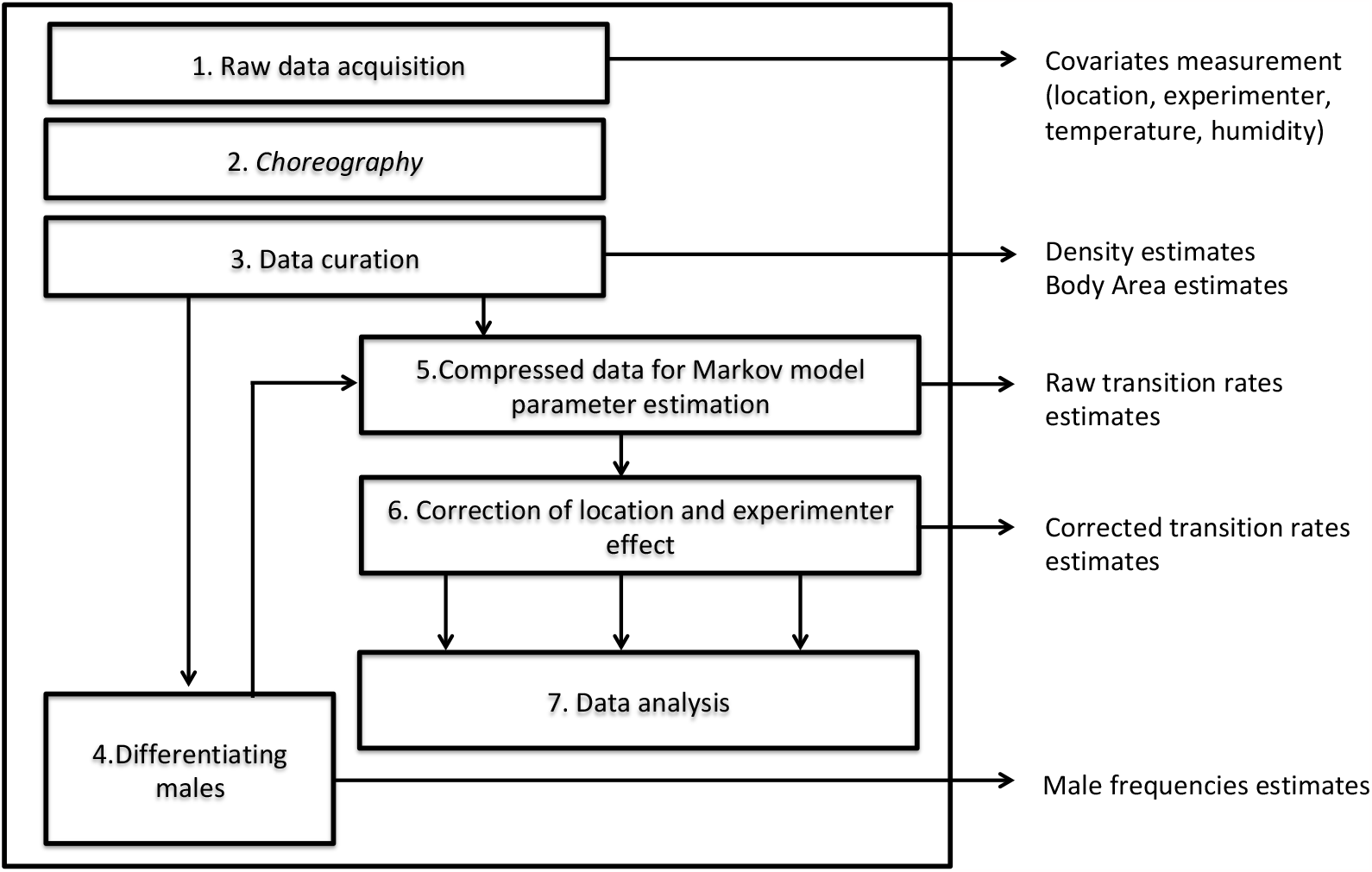
Schematic of data acquisition and analysis pipeline.

**Figure S3:**
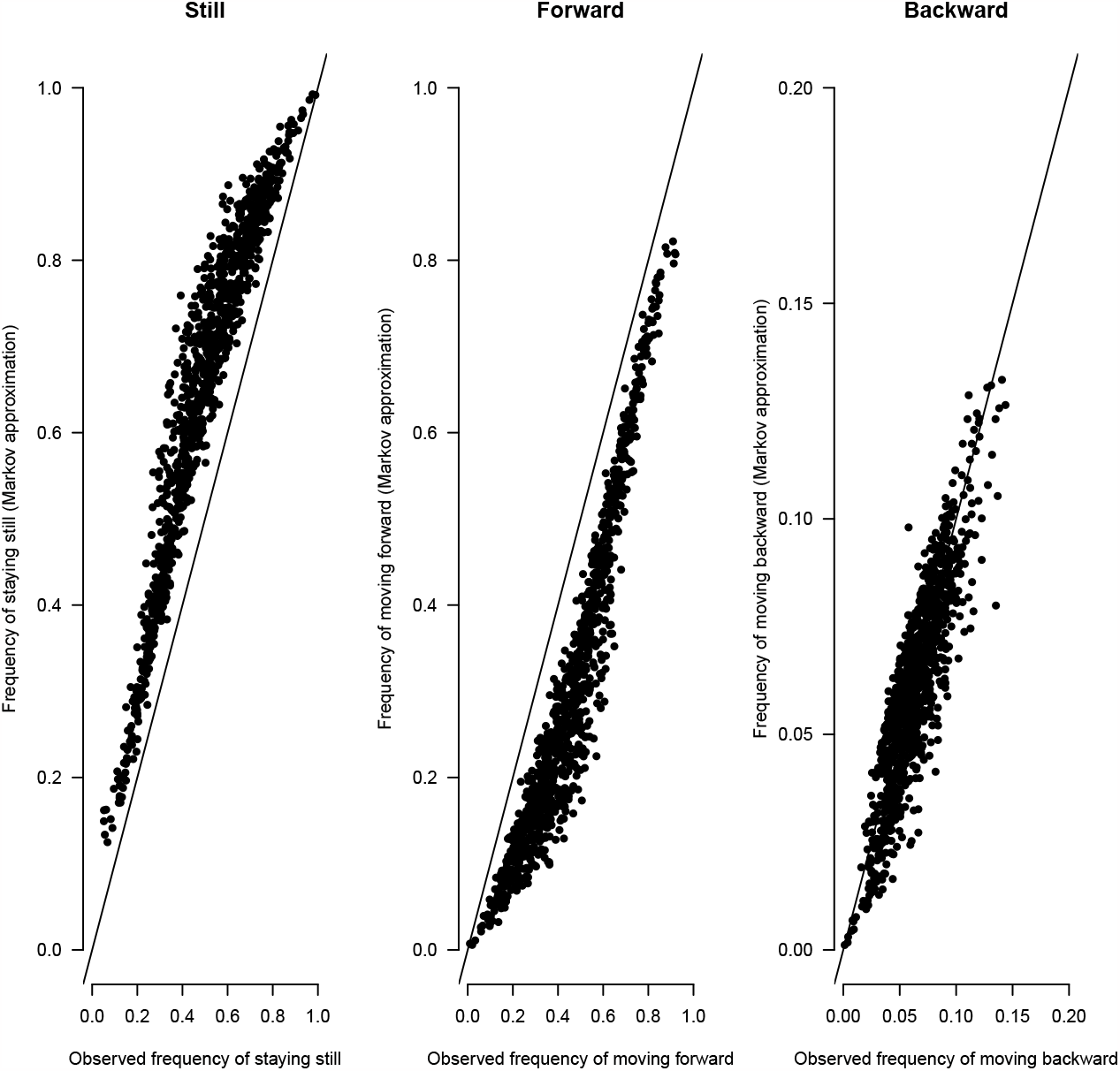
Correlation between the observed frequencies of each of the three movement states and the predicted values from the Markov model. There is a consistent bias in the long term predictions due to violation of the memoryless assumption of the model. Some moving worms tend to remain in this state longer than expected on the long term, that is, they can be briefly interrupted but are more likely to resume movement than predicted.

**Figure S4:**
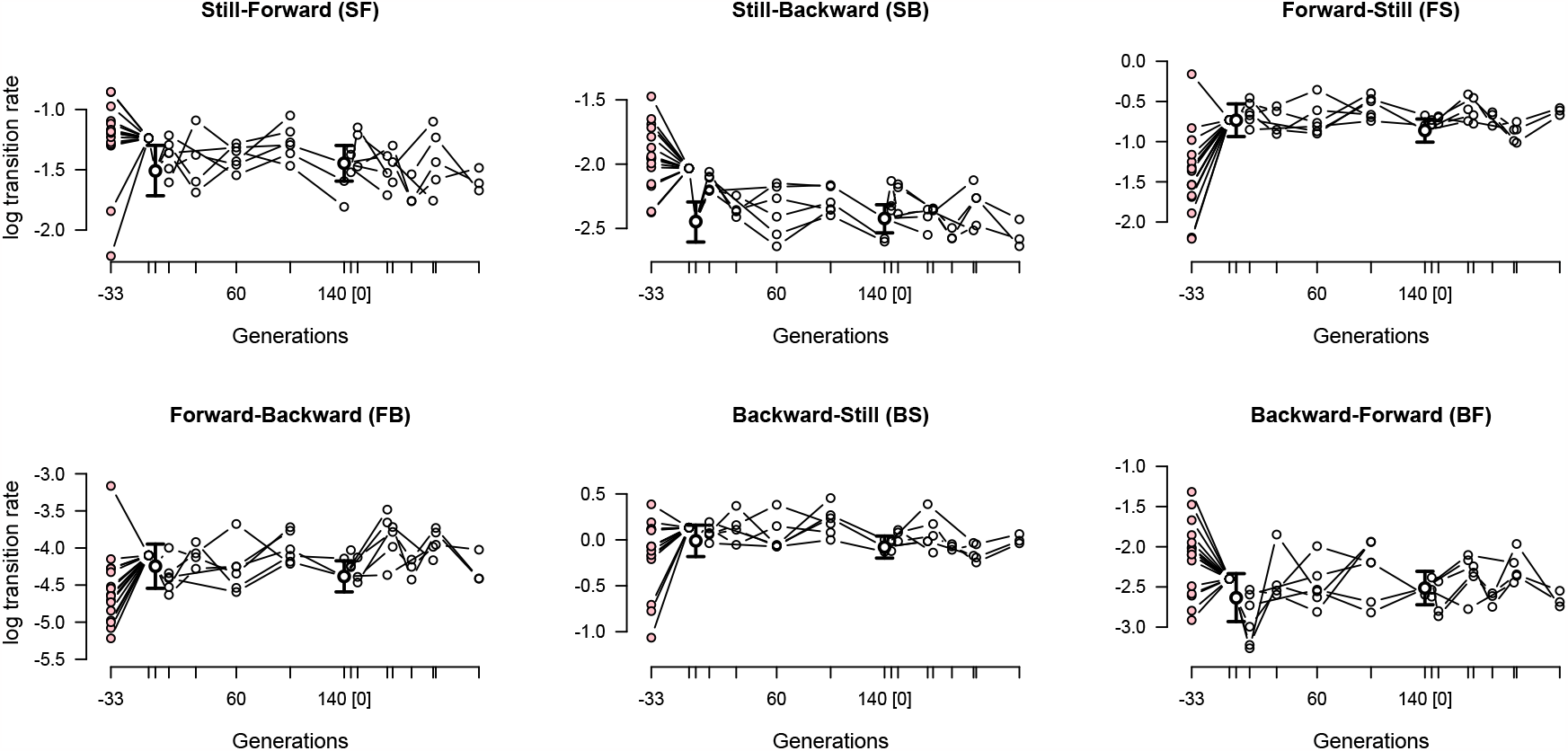
Evolution of mean hermaphrodite transition rates. Each panel shows the evolution of a transition rate in the founders (pink dots) and during experimental evolution (white dots). At the beginning of the domestication and focal stages there was one ancestral population, shown by empty circles with 95% credible intervals, while 3-6 replicate populations were measured at each sampled time point indicated by tick marks.

**Figure S5:**
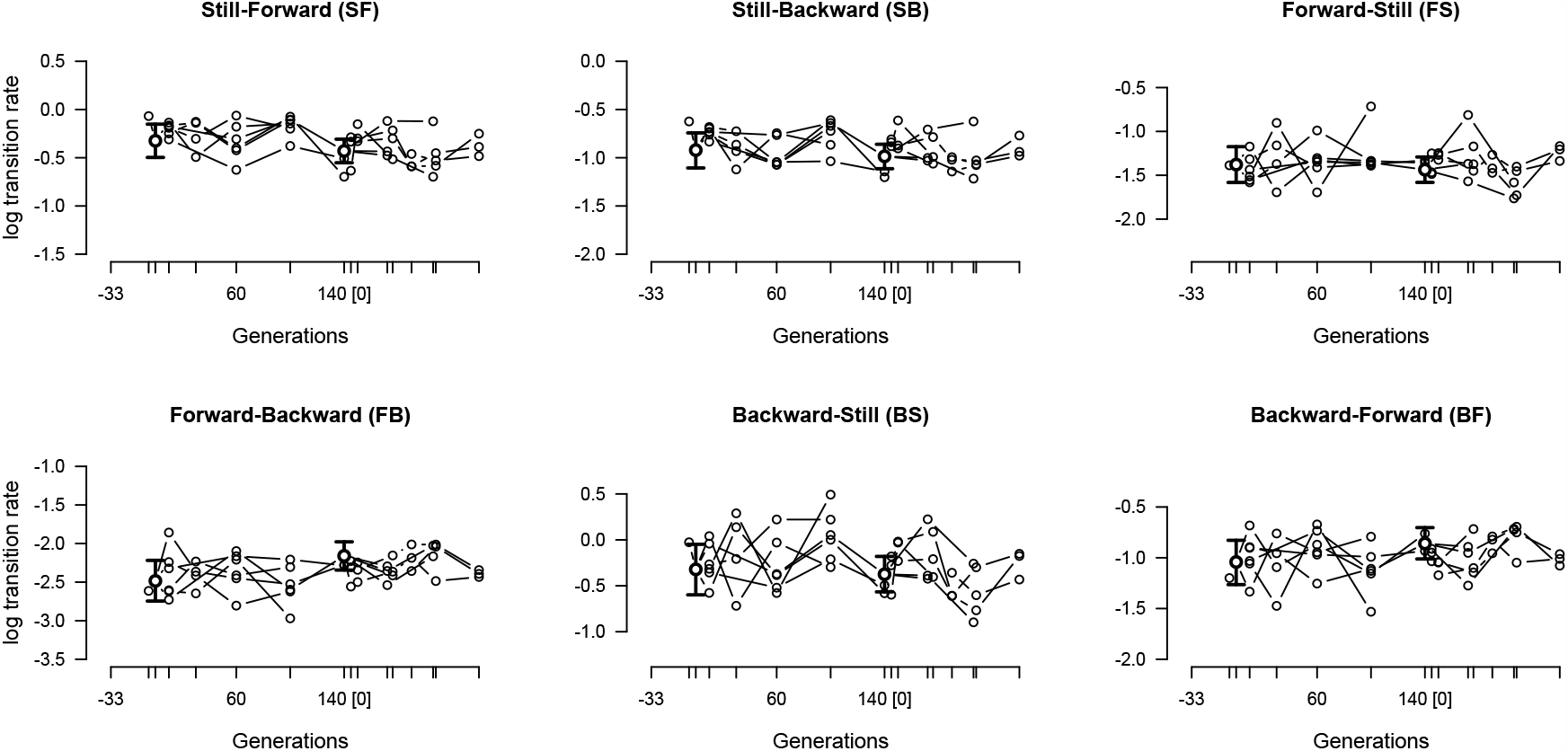
Evolution of mean male transition rates, as in Figure S4. Note that the founder inbred lines do not have any males.

**Figure S6:**
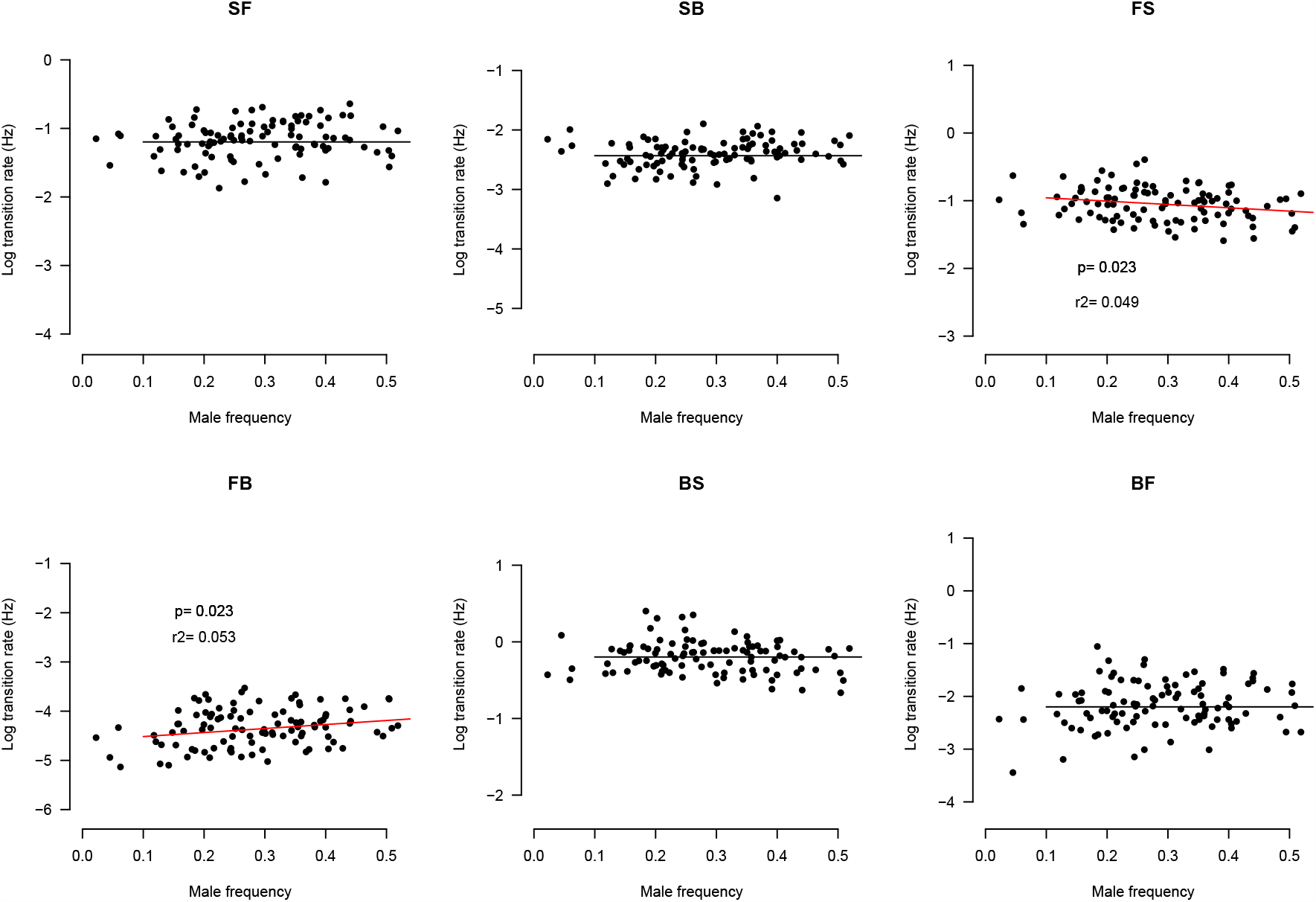
The effects of males on hermaphrodite transition rates in the outbred populations during lab evolution. Each point shows the relation between transition rates and male frequency for each replicate population at a given time point during lab evolution. Red (black) lines show significant (non-significant) linear effects of male frequency on transition rates. For all regression models the coefficient of determination is extremely low (*r*^2^).

**Figure S7:**
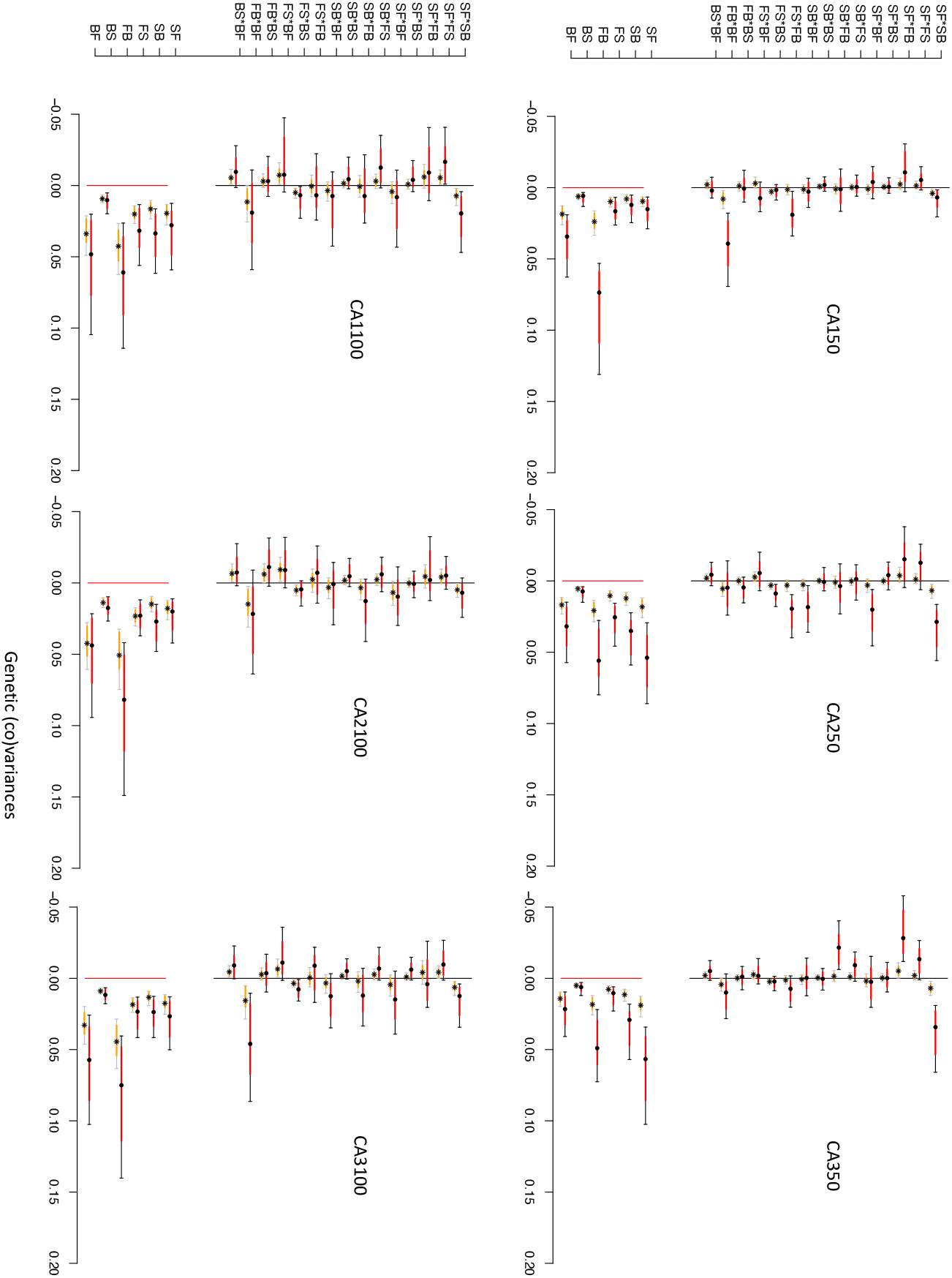
Median genetic covariance and variance estimates of the CA[1-3] populations at generation 50 and generation 100 (dots), as well as random expectations (stars). Intervals are shown with the 95% and 83% credible intervals (black and red, grey and orange). Many of the covariance estimates do not differ from zero (95% interval) and covariance and variance median estimates do not generally differ from a null distribution obtained from random permutations of the phenotypic values (black dots are within grey bars), particularly for generation 100 populations. This is explained by a loss of genetic (co)variances with continued lab evolution after domestication, and not sampling a limited number of lines in each population, see Figure S7).

**Figure S8:**
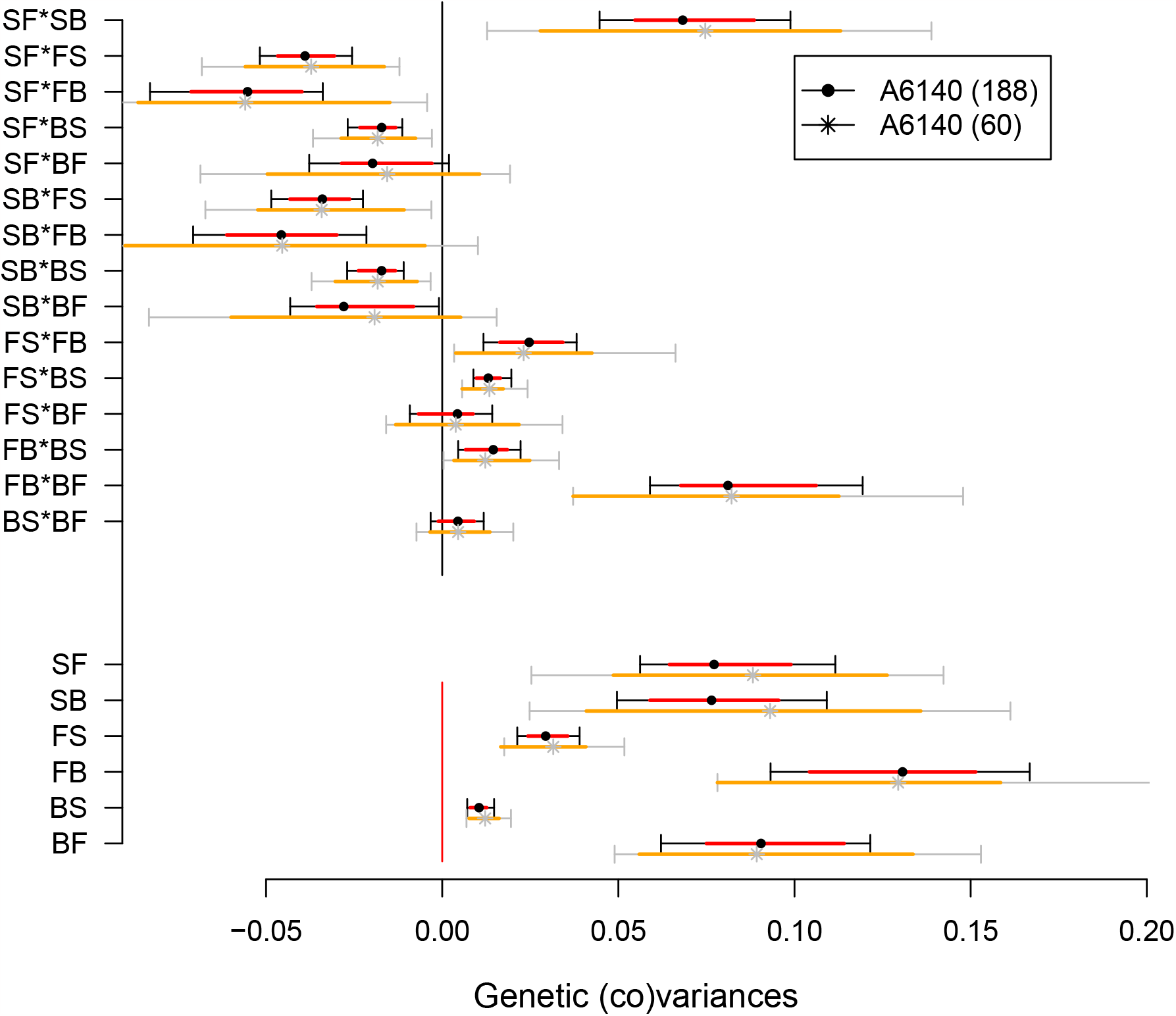
**G**-matrix estimates of the 140-generation domesticated A6140 population. Black and red show the estimated genetic (co)variances using all inbred lines as in main Figure 3. Grey and orange show genetic (co)variances after downsampling to 60 inbred lines, approximately the minimum number of lines phenotyped in the CA[1-3] populations. Median estimates are similar between data sets, though with larger intervals in the subsampled estimates.

**Figure S9:**
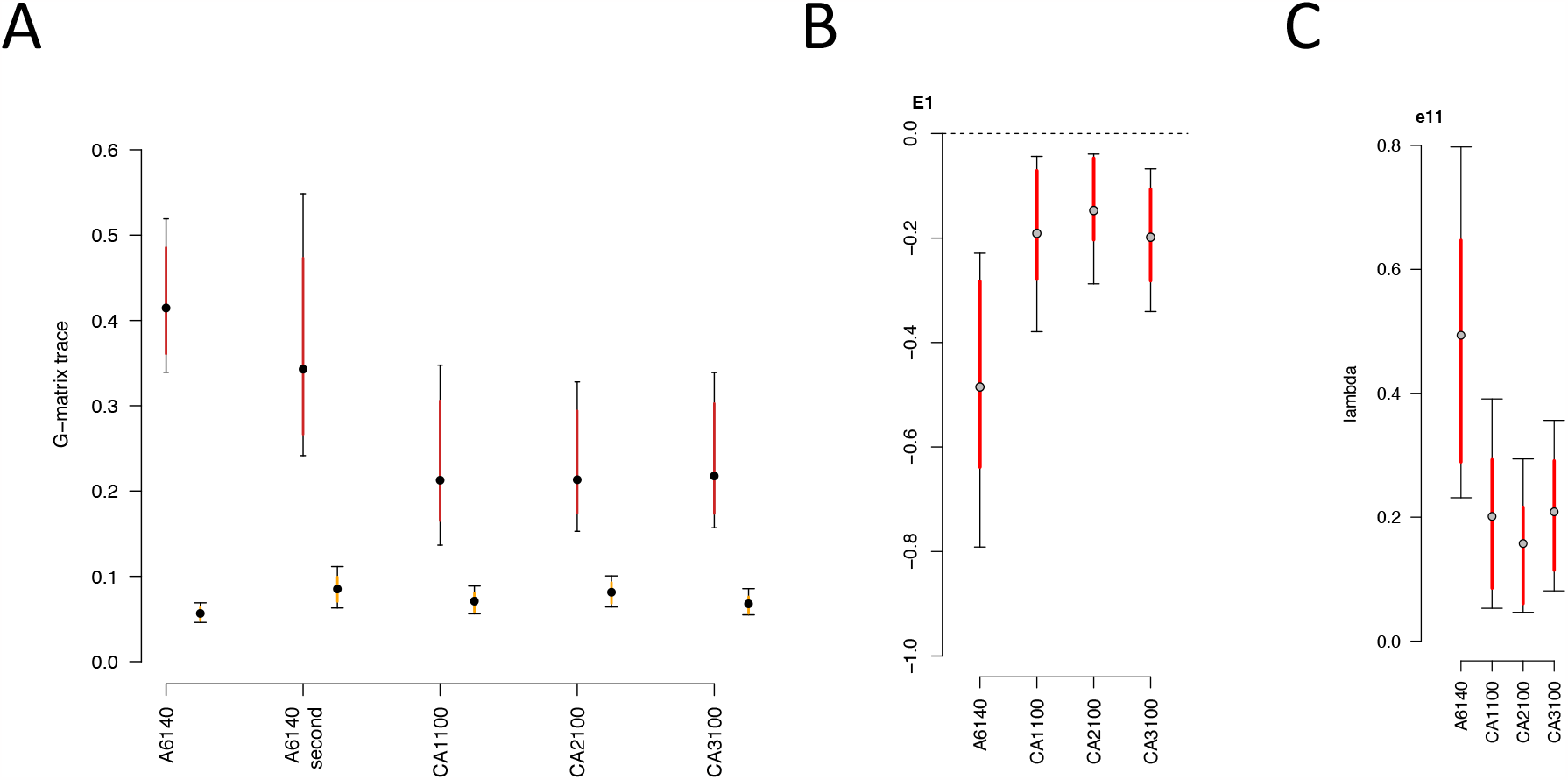
Effect of the common garden assay on genetic (co)variances. **A**. Total amount of genetic variance computed as the sum of the **G**-matrix diagonal elements (trace). The amount of genetic variance in the two A6140 matrices from the two separate common gardens is similar. All observed posterior means differ from the null 95% posterior means (orange). **B**. The coordinates of the **G**-matrices in the space of the first eigentensor when comparing the A6140 and the CA[1-3]100 populations, all computed from the third common garden assay (see Methods). The absolute values of the **G**-matrix coordinates in each eigentensor represent its contribution to the difference between matrices. Coordinates with opposing signs indicate that the matrices contribute in opposing directions. **C**. Contribution of specific trait combinations to coordinated changes among **G**-matrices. Each panel shows the amount of genetic variance in the direction of the greatest variation among **G** (eigenvector of ***E***_**1**_ only). Here, as in panels **A**. and **B**., the results obtained from this second A6140 **G**-matrix are similar to when using the one from the first common garden assay.

**Figure S10:**
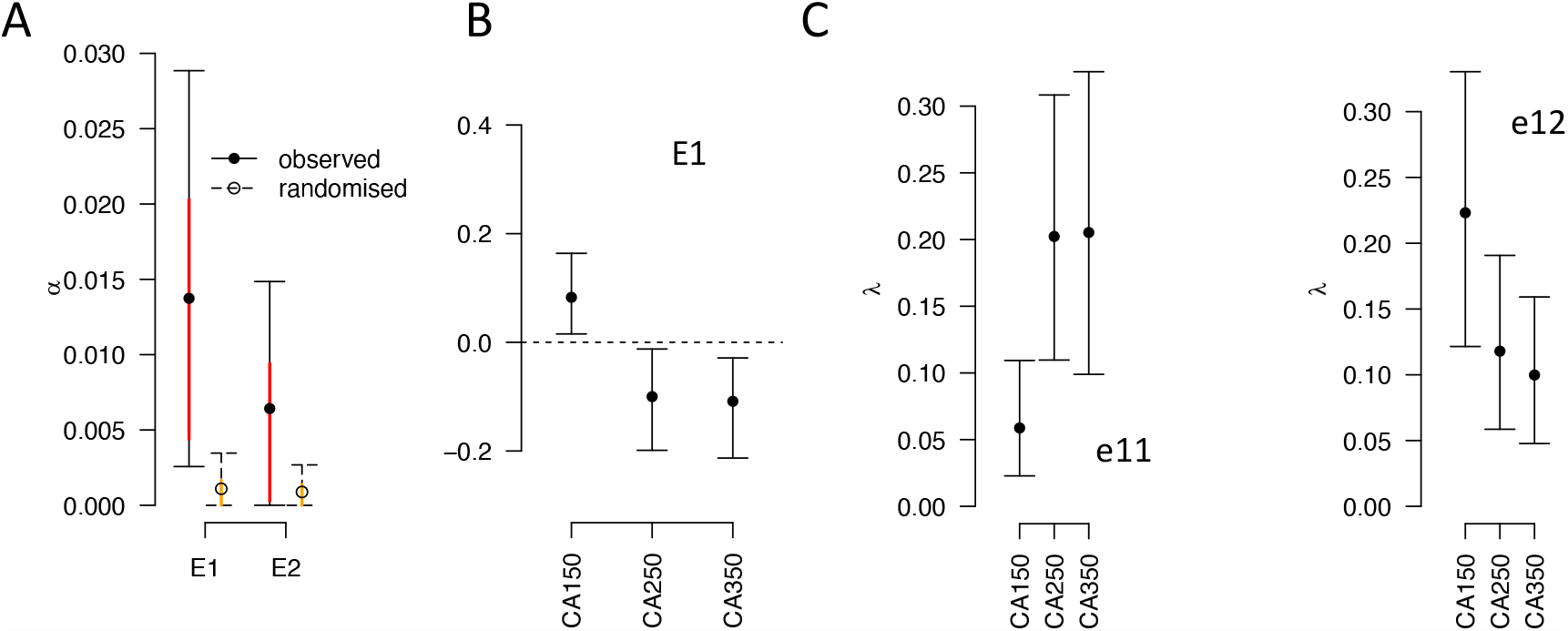
Genetic differentiation. **A**. Eigentensor decomposition of variation among **G**-matrices of the CA[1-3]50 populations. The variance ***α***_***i***_ associated with the *i*th eigentensor ***E***_***i***_ is compared to a null permutation model where variation among matrices is due to sampling (see Methods). Here, only the first eigentensors is different from the null. **B**. The coordinates of the **G**-matrices in the space of the first eigentensor. The absolute values of the matrices coordinates in each eigentensor represent its contribution to the difference between matrices. Coordinates with opposing signs indicate that the matrices contribute in opposing directions. **C**. Contribution of specific trait combinations to coordinated changes among **G**-matrices. Each panel shows the amount of genetic variance in the direction of the greatest variation among **G** (eigenvector of ***E***_**1**_ only).

**Figure S11:**
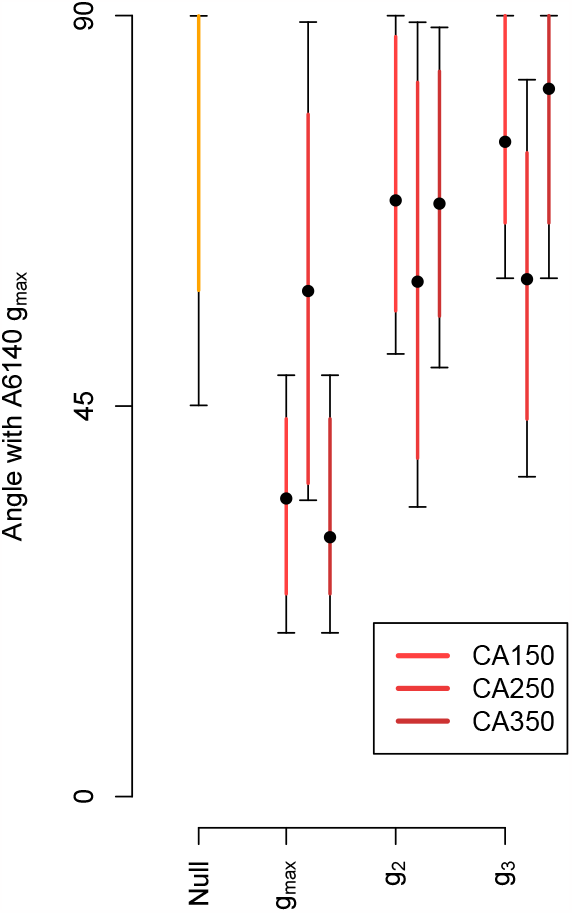
The angle (Θ) between the first three eigenvectors of the evolved **G**-matrix (*g*_*max*_, *g*_1_ and *g*_3_ of the CA[1-3]50 populations) with the A6140 *g*_*max*_. Θ differs from the random expectations in CA150 and CA350 but not in CA250 showing genetic differentiation. Dots show the mean estimate with bars the 83% and 95% credible interval of the posterior **G**-matrix distribution. The null expectation was obtained by computing the angle between pairs of random vectors sampled from a uniform distribution.

**Figure S12:**
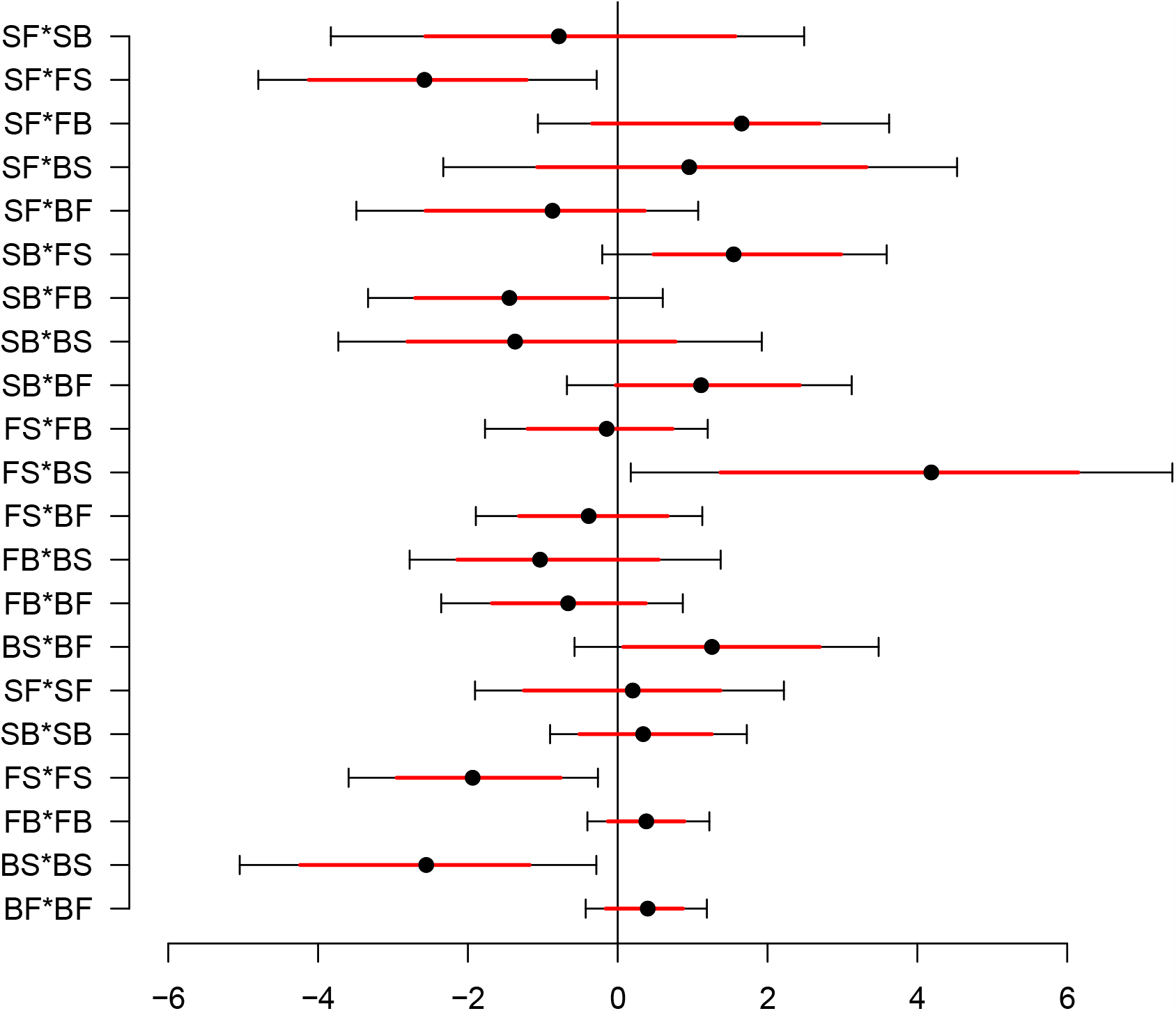
Quadratic selection coefficients. The partial regression coefficients of fertility on trasition rates estimated by Bayesian inference. Each row shows the mode (dot), and 83% and 95% credible intervals (red bar and line bars, respectively) of the posterior distributions. The top 15 rows show coefficients of correlated selection between two transition rates, the bottom 6 rows show coefficients of stabilizing or disruptive selection on each transition rate.

**Figure S13:**
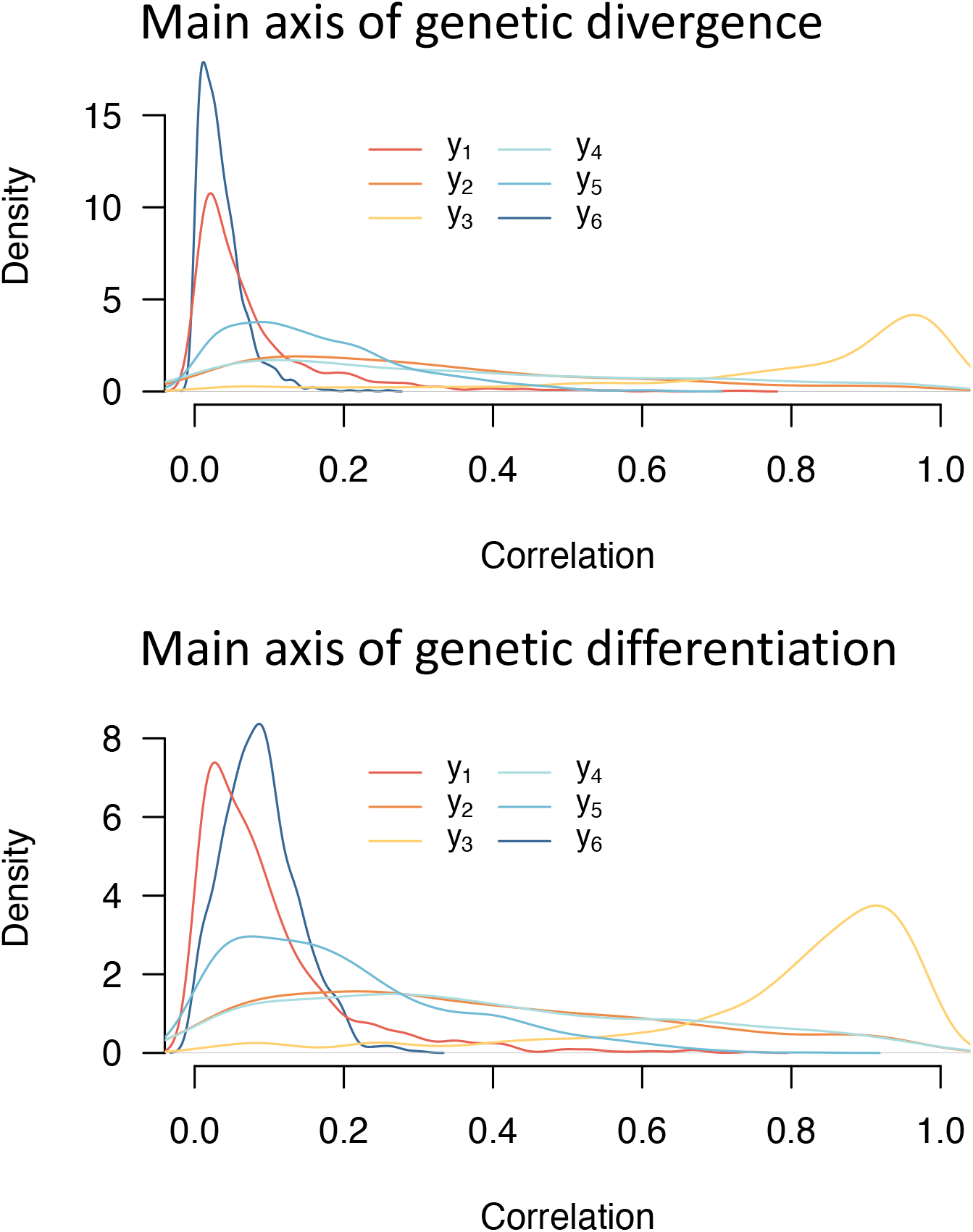
Alignment of **G**-matrices divergence and differentiation with the quadratic selection surface. Shown is the density distributions of Pearson product moment correlations between the first eigenvector *e*_11_ of **E1** measured for divergence (between A6140 and CA[1-3]100, top panel) and for genetic differenciation (measured among CA[1-3]50, bottom panel). The density distributions are obtained from 1000 sampling in the posterior distribution of the *γ* matrix.

